# Phasor histone FLIM-FRET microscopy quantifies spatiotemporal rearrangement of chromatin architecture during the DNA damage response

**DOI:** 10.1101/419523

**Authors:** Jieqiong Lou, Lorenzo Scipioni, Belinda K. Wright, Tara K. Bartolec, Jessie Zhang, V. Pragathi Masamsetti, Katharina Gaus, Enrico Gratton, Anthony J. Cesare, Elizabeth Hinde

## Abstract

To investigate how chromatin architecture is spatiotemporally organised at a double strand break (DSB) repair locus, we established a biophysical method to quantify chromatin compaction at the nucleosome level during the DNA damage response (DDR). The method is based on phasor image correlation spectroscopy (ICS) of histone FLIM-FRET microscopy data acquired in live cells co-expressing H2B-eGFP and H2B-mCherry. This multiplexed approach generates spatiotemporal maps of nuclear-wide chromatin compaction that when coupled with laser micro-irradiation induced DSBs, quantify the size, stability, and spacing between compact chromatin foci throughout the DDR. Using this technology, we identify that ATM and RNF8 regulate rapid chromatin decompaction at DSBs and formation of a compact chromatin ring surrounding the repair locus. This chromatin architecture serves to demarcate the repair locus from the surrounding nuclear environment and modulate 53BP1 mobility.

**SIGNIFICANCE STATEMENT:** Chromatin dynamics play a central role in the DNA damage response (DDR). A long-standing obstacle in the DDR field was the lack of technology capable of visualising chromatin dynamics at double strand break (DSB) sites. Here we describe novel biophysical methods that quantify spatiotemporal chromatin compaction dynamics in living cells. Using these novel tools, we identify how chromatin architecture is reorganised at a DSB locus to enable repair factor access and demarcate the lesion from the surrounding nuclear environment. Further, we identify novel regulatory roles for key DDR enzymes in this process. Finally, we demonstrate method utility with physical, pharmacological and genetic manipulation of the chromatin environment, identifying method potential for use in future studies of chromatin biology.

## INTRODUCTION

The DNA damage response (DDR) is constantly scanning chromatin for genomic lesions (1-5). In the event DNA damage is detected, the DDR signals the lesion’s presence, mediates downstream repair, and arrests cell growth until chromatin restoration is complete (6-8). Following induction of a DSB, the ataxia-telangiextasia mutated (ATM) kinase localises to the lesion to regulate cell cycle arrest and signal recruitment of repair factors to the damaged chromatin (2, 9-12). This includes the RNF8 E3 ubiquitin-protein ligase (11, 13-16), which initiates ubiquitin signalling at the lesion that is essential for recruitment of additional repair factors such as 53BP1 (16-18). In addition, the DDR also recruits proteins to the repair locus that contribute to chromatin decompaction or nucleosome eviction (3, 13, 19). Paradoxically, factors mediating chromatin compaction, such as heterochromatin protein 1, also localise to DSBs to facilitate repair (20-23). Currently it remains unclear how these contrasting chromatin dynamics coincide at a DSB repair locus, and how this impacts chromatin macrostructure elsewhere in the nucleus. Interrogating DDR-dependent chromatin dynamics requires a technology capable of simultaneously probing chromatin structure at DNA lesions and throughout the entire nucleus, during the entirety of the DNA repair process.

The most basic feature of chromatin structure is the nanometre spacing (2-50 nm) between nucleosomes (24-26). Given that the scale of this structural subunit is below the diffraction limit of optical and most super resolution imaging modalities (27), the rearrangements in chromatin structure during the DDR are rendered ‘invisible’. To get around this technical hurdle, fluorescence recovery after photobleaching (FRAP) (24), single-particle tracking (SPT) (28, 29), and fluorescence correlation spectroscopy (FCS) (30, 31) have been employed to measure molecular diffusion within the chromatin landscape and indirectly probe chromatin nanostructure (32-35). These methods, however, do not directly observe chromatin compaction during DNA repair. What is needed to visualise changes in chromatin architecture is an experimental means to directly observe chromatin organisation in live cells, at the level of nucleosome packaging, throughout the entirety of the nuclear volume, and for the duration of the DDR.

To bridge this experimental gap, here we establish the phasor approach to fluorescence lifetime imaging microscopy (FLIM) of Förster resonance energy transfer (FRET) interaction between fluorescently labelled histones. FRET is exquisitely sensitive to fluorophore proximity and can report on changes in nucleosome spacing when the core histone H2B is individually tagged to donor and acceptor fluorophores (36-38). By coupling this readout of chromatin structure with DSB induction using near infrared (NIR) laser micro-irradiation (35, 39), we are able to directly measure nanometre changes in chromatin compaction at a DNA damage site versus globally throughout the nucleus. Furthermore, because phasor analysis enables fit free quantification of spectroscopic properties recorded in each pixel of an image (40-42), FRET maps of chromatin foci are amenable to image correlation spectroscopy (ICS) (43). ICS is a general approach to quantitate fluorophore distribution within an image (44-47) and phasor transformation of local image correlation functions (PLICS) uncovers heterogeneity within fluorophore number, size and spatial distribution (47). Therefore, in the context of histone FRET, phasor-based ICS enables nuclear wide chromatin organisation to be spatiotemporally characterised at the level of nucleosome proximity and in live cells.

In this study we use phasor-ICS analysis of histone FLIM-FRET in HeLa cells stably co-expressing low levels of H2B-eGFP and H2B-mCherry (Hela^H2B-2FP^) to investigate how chromatin architecture is organised at DSBs by the DDR. We find that following DSB induction with a NIR laser, ATM and RNF8 regulate chromatin decompaction at the damage site coupled with formation of a surrounding stable ring of compacted chromatin. Correlation of chromatin architecture with measurements of 53BP1 accumulation and mobility reveal that DNA repair activity is restrained to the decompacted chromatin environment at the lesion, and that the surrounding compact chromatin ring demarcates the DNA repair locus from the nuclear environment whilst controlling 53BP1 mobility. Collectively, these data reveal how chromatin architecture is organised by the DDR and demonstrate that phasor ICS analysis of histone FLIM-FRET is amenable to the study of local and global chromatin dynamics under conditions of physical, pharmacological or genetic alteration.

## RESULTS

### Phasor analysis of chromatin compaction in live cells using FLIM-FRET microscopy

To assay chromatin compaction at the level of nucleosome proximity, we generated HeLa^H2B-2FP^ cells stably co-expressing histone H2B-eGFP and histone H2B-mCherry (Fig 1a, Supplementary Fig. 1). Tagged histone cDNAs were transduced via retrovectors, selected for transgene expression, and cells sorted for low eGFP and mCherry expression to minimise the impact of exogenous transgene expression on cellular function. Stable co-expression of both histones into chromatin was confirmed by live cell imaging (Supplementary Fig. 1a), which also revealed no increase in underlying genome instability or mitotic abnormalities in HeLa^H2B-2FP^ (Supplementary Fig. 1b-f).

**Figure 1.**
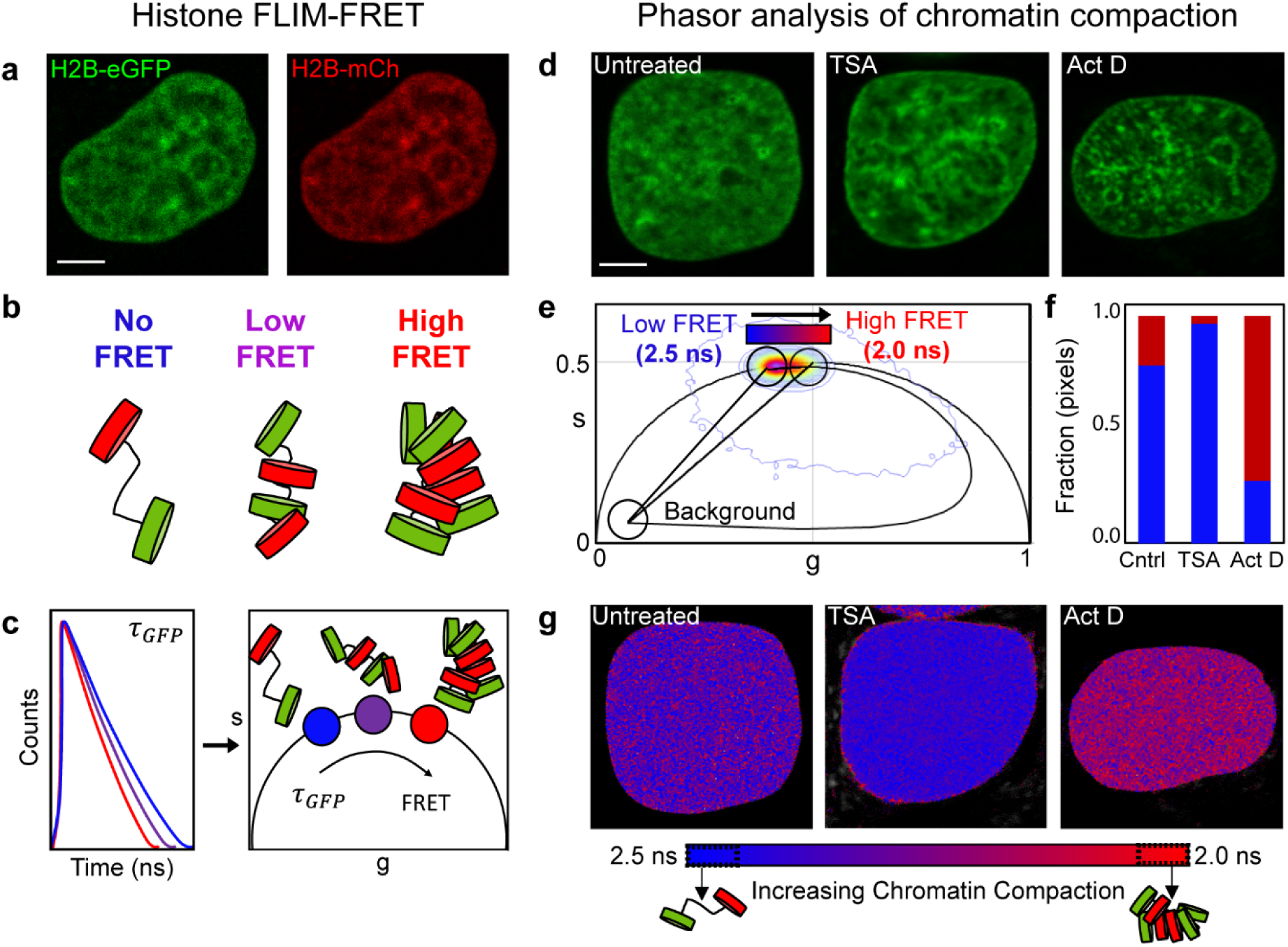
Phasor approach to FLIM-FRET analysis of chromatin compaction. (a) HeLa^H2B-2FP^ nucleus co-expressing H2B-eGFP and H2B-mCherry. (b) Graphical depiction of how increasing nucleosome proximity leads to increased FRET between fluorescent histones. (c) Graphical depiction of phasor transformation of HeLa^H2B-2FP^ FLIM-FRET data. Left: fluorescence lifetime of H2B-eGFP reports on the degree of FRET interaction in each pixel. Each line represents the fluorescent lifetime from a different pixel. Right: These data when phasor transformed give rise to phasor coordinates (s, g). The phasor trajectory is right shifted to shorter fluorescent lifetimes depending on FRET efficiency. In HeLa^H2B-2FP^ decreasing lifetime and increasing FRET corresponds to more compact chromatin. (d) Untreated and Trichostatin A (TSA) or Actinomycin D (Act D) treated HeLa^H2B-2FP^ nuclei, shown in the H2B-eGFP channel. (e) Combined phasor distribution of H2B-eGFP fluorescence lifetime from all conditions shown in (d) defines the range of FRET efficiency in HeLa^H2B-2FP^. (f) Fraction of pixels in a compact (red) versus open (blue) chromatin state in TSA and Act D treated cells (g) Pseudo-coloured chromatin compaction maps of the cells in (d) according to palette defined in the phasor plot data in (e). Scale bar equals 5 μm.

Using HeLa^H2B-2FP^ we established a workflow of FLIM-FRET data acquisition and phasor lifetime analysis to quantify the local chromatin compaction status within each pixel of a HeLa^H2B-2FP^ nucleus. FRET detects molecular interaction within 1-10 nm and the efficiency of energy transfer between a donor and acceptor molecule is exquisitely sensitive to distance (48, 49). Increasing FRET efficiency in HeLa^H2B-2FP^ corresponds to a reduced spacing between nucleosomes containing H2B-eGFP (donor) and nucleosomes containing H2B-mCherry (acceptor) (graphically depicted in Fig. 1b). FRET efficiency can be quantified by measurement of the fluorescence lifetime of H2B-eGFP, as this property is increasingly quenched upon closer interaction with H2B-mCherry. Therefore, in HeLa^H2B-2FP^ a reduced H2B-eGFP fluorescence lifetime corresponds with increased chromatin compaction as nucleosomes move into closer proximity.

To measure the fluorescence lifetime of H2B-eGFP in each pixel of an HeLa^H2B-2FP^ nuclei, we used 2-photon excitation on a laser-scanning confocal microscope equipped with time resolved detection. To translate fluorescence lifetime into FRET efficiency, we employed phasor analysis (graphically depicted in Fig. 1c). Phasor analysis transforms the fluorescence lifetime recorded in each pixel of a FLIM image into a two-dimensional coordinate system (g, s) (40, 41). When these coordinates are placed in a single phasor plot, the phasor coordinates derived from each pixel give rise to a series of clusters which fingerprint all the measured FRET efficiencies within a FLIM image. Specifically, pixels in a HeLa^H2B-2FP^ nucleus which contain compacted chromatin, and therefore exhibit a reduced H2B-eGFP fluorescence lifetime, have an increasingly right shifted phasor value (Fig. 1c).

The phasor method enables quantitation of FRET in each pixel of a FLIM image and spatially maps the different chromatin states throughout a HeLa^H2B-2FP^ nucleus. To establish the dynamic range of chromatin compaction states detectable in our experimental system, we treated HeLa^H2B-2FP^ with the histone deacetylase inhibitor Trichostatin A to loosen chromatin structure (32), or the transcriptional inhibitor Actinomycin D to compact the chromatin network (50) (Fig. 1d, Supplementary Fig. 2). Phasor coordinates recorded from the FLIM data in untreated and treated cells were collated into a single phasor plot (Fig. 1e) and found to detect varying degrees of chromatin compaction between a fluorescence lifetime of 2.5 ns (TSA treatment, low compaction) and 2.0 ns (Actinomycin D treatment, high compaction). This lifetime change corresponds to a 0 to 21% change in FRET efficiency and enables us to respectively define the phasor location of ‘open’ versus ‘compacted’ chromatin. These parameters were used throughout this study to define and pseudo-colour heterogeneous chromatin compaction states recorded in each pixel of a FLIM acquisition (Fig. 1f, g).

### Chromatin compaction is spatiotemporally controlled during the DNA damage response

To investigate chromatin compaction during the DDR in live HeLa^H2B-2FP^ cells we coupled phasor FLIM-FRET with NIR laser micro-irradiation. This multiplexed approach enabled induction of DSBs within a defined region of interest (ROI) and measurement of local versus global rearrangements in chromatin structure as a function of time. NIR laser micro-irradiation can be tuned to induce different types of DNA damage by modulating the laser frequency, wavelength or power (19, 39, 51, 52). To determine conditions that generate DSBs in HeLa^H2B-2FP^, we identified laser settings that induced lesions which recruited eGFP-53BP1 or Ku-GFP to the irradiated ROI in transiently transfected parental HeLa cells (Supplementary Fig. 3) (35). These settings were used for the remainder of the experimentation described below.

For all experiments we measured fluorescence intensity (Fig. 2a) and lifetime (Fig. 2b) of H2B-eGFP in each pixel of a HeLa^H2B-2FP^ nucleus immediately before DSB induction, immediately after DSB induction (T=0) and at regular intervals for the following six hours. Laser micro-irradiation can photo-bleach H2B-eGFP at the damage site. However, given that the fluorescence lifetime of H2B-eGFP is independent of concentration, this artefact does not impact on FRET determination of chromatin compaction (49, 53). From the FLIM maps derived during the DDR (Fig. 2b) we observed that immediately after NIR laser micro-irradiation, chromatin compaction is induced at the damage site (red pixels). Then, within the following hour a compacted chromatin architecture propagates outward to form a border surrounding a central region of open chromatin at the DNA lesion (Fig. 2c).

**Figure 2.**
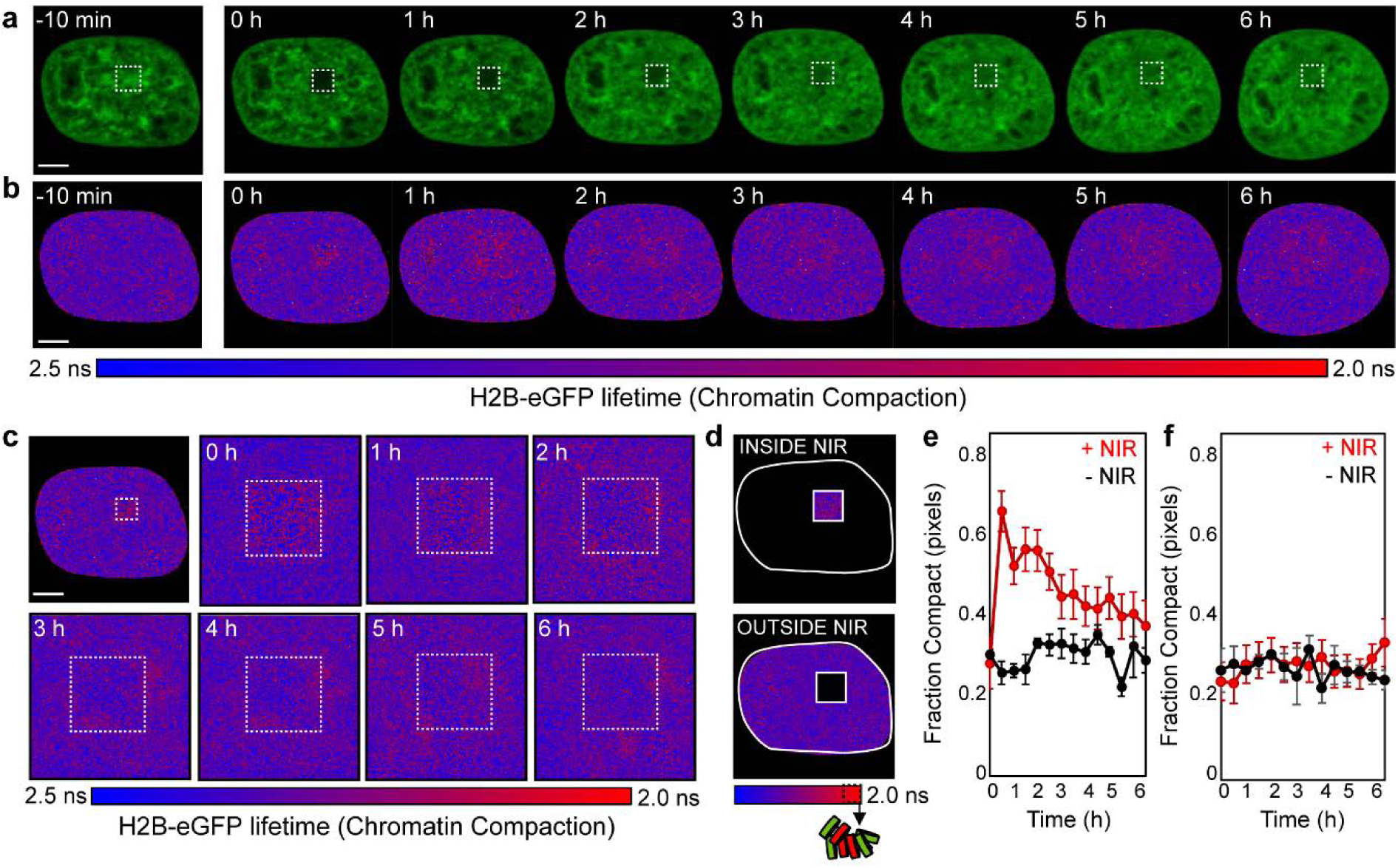
FLIM-FRET analysis of chromatin compaction reveals chromatin architectural changes during the DDR. (a, b) Time series of H2B-eGFP fluorescence intensity images (a) and lifetime maps (b) acquired in a HeLa^H2B-2FP^ before and at hourly intervals after NIR irradiation. The white square indicates the NIR laser treated locus. (c) Digital enlargement of the DNA damage site selected in (b) and the corresponding time series of lifetime maps within this region of interest. (d) Masks selected for analysis of the number of pixels in a compacted (high FRET) versus non-compacted (low FRET) state at the DNA damage site versus outside this ROI. (e, f) Fraction of pixels within (e) and outside (f) of the NIR irradiated ROI that are in a compacted state during the DNA damage response (red curve) versus an unperturbed cell (black curve) (N=10 cells, mean ± s.e.m.). Scale bar equals 5 μm.

To quantify the temporal progression of chromatin compaction at the DNA damage site, and any potential changes in global chromatin network organisation, we next quantified the fraction of pixels in a compacted state inside and outside of the NIR treated ROI (Fig. 2d). This revealed: (1) the fraction of pixels with compact chromatin inside the NIR treated locus sharply increased in the first 30 min after DSB induction and persisted for up to 3 hours (Fig. 2e), and (2) there was no significant change in the percentage of pixels with compact chromatin in the nuclear wide undamaged chromatin network outside of the NIR treated ROI (Fig. 2f). While informative, this analysis masks spatial rearrangement in compact chromatin foci localisation at the DNA damage site. Interrogation of the spatiotemporal dynamics that underlie rearrangement of compact chromatin foci into a local border surrounding an opened DNA lesion (Fig. 2c) required development of analytical tools capable of quantifying the temporal stability and spatial reorganisation of compact chromatin in a single cell over time.

### Image correlation analysis of FRET quantifies global chromatin network organisation

To quantify the stability, size, and spacing of compact chromatin foci we first extracted the coordinates of pixels in a high FRET state within a FLIM acquisition (Fig. 3a-b). This derived a spatial map of compact chromatin localisation (τH2B-eGFP= 2.0 ns, Fig. 3c). To identify which of these compact chromatin foci persist in a cell over time, we then developed a novel image correlation analysis method termed longevity analysis (Fig. 3d-f). In longevity analysis a moving average is applied across a time series of chromatin compaction maps derived from FLIM-FRET data (Fig. 3d) to produce a longevity map (graphically presented in Fig. 3e). Because the source localisation images are binary, the amplitude recorded in each pixel directly reports how long a structure persists in time. Longevity analysis therefore enables us to extract and spatially map stable versus dynamic compact chromatin foci (Fig. 3f, left), and provides a quantitative read-out of chromatin network dynamics that can be tracked throughout the nucleus for the entirety of the DDR (Fig. 3f, right).

**Figure 3:**
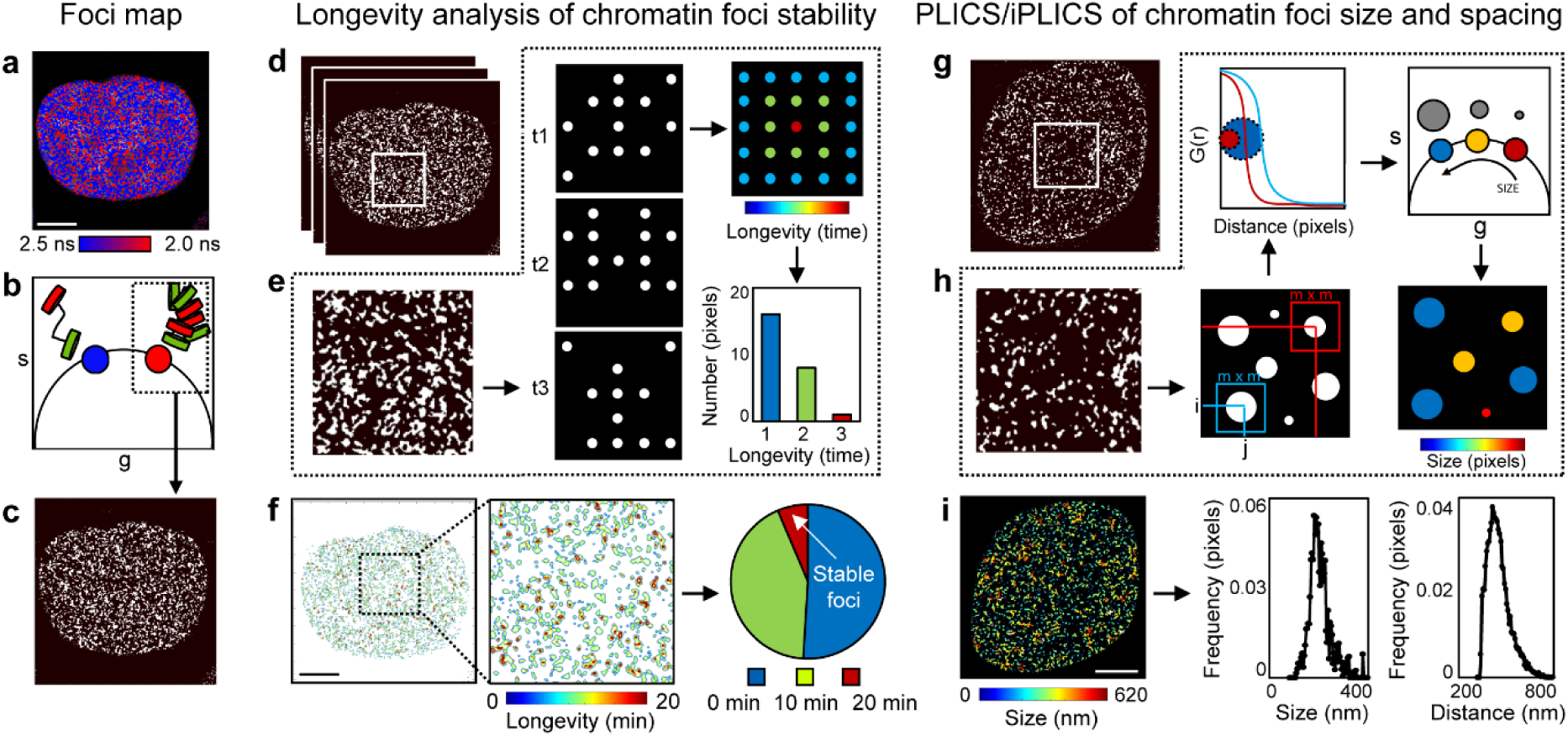
Longevity and PLICS/iPLICs analysis measures nuclear wide compact chromatin stability, size, and spacing. *Foci map*: (a) FLIM-FRET map from an unirradiated HeLa^H^2^B-^2^FP^ nucleus. (b, c) Pixel coordinates of high FRET state (τH2B-eGFP = 2.0 ns) can be extracted from phasor plot (b) to produce a localisation map of ‘compact’ chromatin (c). *Longevity analysis*: (d) Localisation map of compacted chromatin from the HeLa^H2B-2FP^ cell in (a) at 0, 10 and 20 min (as detected by FRET). (e) Schematic of longevity analysis: averaging three binary images gives rise to a heat map of pixel longevity, which contains structural information that is not evident in the source images and can be used to quantify the stability of detected structures. (f) Longevity map of the compacted chromatin foci detected and tracked in (d), with digital enlargement shown for a region of interest that contains foci present for 10 min (green pixels) and 20 min (red pixels). The fraction of foci persistence across the time course are calculated as a measure of overall chromatin network stability. *PLICS analysis:* (g) A localisation map of compacted chromatin from an unirradiated HeLa^H2B-2FP^ nucleus as detected by FLIM-FRET. (h) Schematic of PLICS analysis: in a binary image showing different sized structures we can calculate localised two-dimensional spatial correlation functions using a m × m matrix and by collapsing them into one-dimensional correlation profiles. The resulting decay is characteristic of the size of a structure within each m × m matrix. By transforming each decay into phasor coordinates (g, s), we can graphically pseudo-colour each pixel according to size. (i) Size map of compacted chromatin foci detected in (g) and PLICS/iPLICS analysis of the average size (305 nm) and spacing (485 nm). Scale bars equal 5 μm.

Longevity analysis identifies when and where compact chromatin foci undergo a rearrangement. However, it does not reveal how compact chromatin spatially redistributes within a given time window. To visualise this, we established an additional method based on a phasor transformation of local image correlation functions (also termed “PLICS”) (47) to uncover the size and spacing between compact chromatin foci (Fig. 3g-i). PLICS analysis starts with a localised spatial autocorrelation analysis within square (m × m) matrices that surround each pixel (*i,j*) in a derived chromatin localisation map (graphically presented in Fig. 3h). This gives rise to a series of two-dimensional autocorrelation functions (ACFs) that when collapsed into a one-dimensional decay, describe the size of the objects centred at that pixel. The local ACF in each pixel is then transformed into a set of phasor coordinates (g, s), which when placed in a single phasor plot identify the different sized compact chromatin structures (top row, Fig. 3h). The phasor coordinates corresponding to different sized compact chromatin foci are then pseudo-coloured and spatially mapped back to the source image (bottom right, Fig. 3h).

Next, we extended PLICS analysis to a novel “inverse PLICS (iPLICS)” approach that enabled us to identify the spacing between compact chromatin clusters (Supplementary Fig. 4a). Here the negative of a chromatin compaction map is first derived. Then, when these data are analysed through the PLICS workflow, we calculate localised two-dimensional spatial correlation functions that when collapsed into a one-dimensional ACF describe the characteristic spacing between structures within a square (m × m) matrix. Transforming each decay into phasor coordinates (g, s) enables us to identify spacing between compact chromatin foci and pseudo-colour the source image. To demonstrate the feasibility of PLICS and iPLICS, we simulated an image containing circular binary structures with a diameter of 3 pixels that were evenly spaced by 8 pixels (Supplementary Fig. 4b), and correctly recovered the size and spatial distribution of these structures (Supplementary Fig. 4c). Thus, when PLICS and iPLICS were applied to the unirradiated HeLa^H2B-2FP^ nucleus shown in Figure 3g, we were able to detect compact chromatin foci between 0-620 nm (Fig. 3i, left) with an average compact chromatin foci size of 305 nm and spacing of 485 nm between structures (Fig 3i, right). Collectively these parameters characterise the nuclear wide spatial organisation of local chromatin compaction.

### Uniform induction and heterogenous resolution of chromatin organisation in the DDR

We applied the longevity, PLICS, and iPLICS workflow to phasor FLIM-FRET datasets from NIR irradiated HeLa^H2B-2FP^ cells (Fig 4a). Longevity analysis was performed using a moving average of 10 min during the first hour after NIR irradiation (Fig. 4b), and every hour for six hours post irradiation (Supplementary Fig. 5a,b). This revealed the strong induction of a stable and compact chromatin border around the damage site, particularly 10-30 min post-irradiation (Fig 4b). Plotting stable compact chromatin foci revealed the chromatin network is more dynamic during the first 30 min after DSB induction, before stabilising by 60 min (Fig. 4c). Notably, while a stable compact chromatin border was uniformly generated in all cells analysed within the first 30 min following irradiation, we observed inter-cell heterogeneity in resolution of this structure over the course of six hours (Supplementary Fig. 5c).

**Figure 4:**
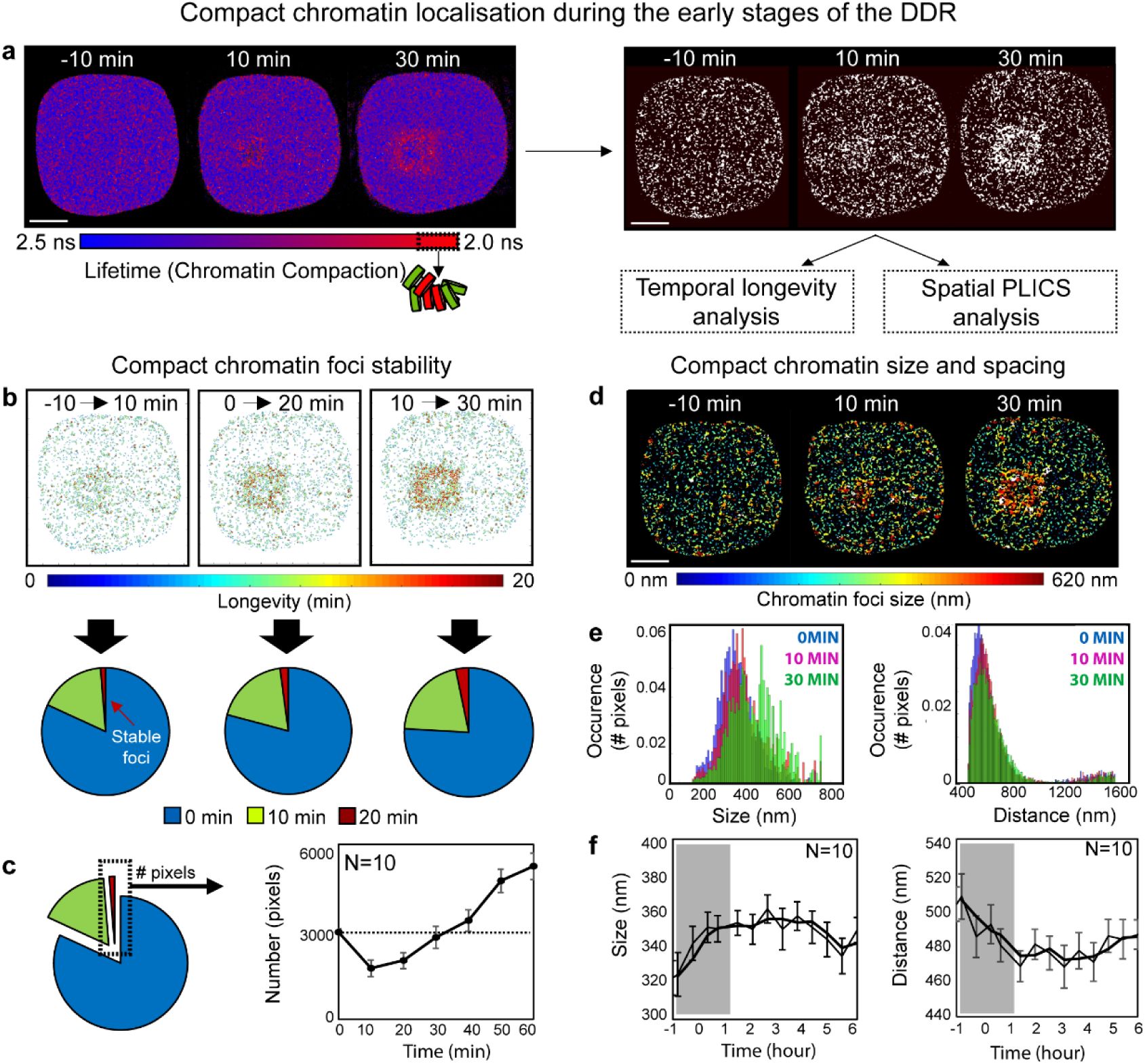
A compact chromatin border forms around the lesion site within the first 30 min of the DDR. (a) FLIM-FRET maps and their derived compact chromatin maps acquired in a HeLa^H2B-2FP^ cell during the first 30 min after NIR irradiation. (b) Quantification of longevity maps of compact chromatin foci taken at 10 min intervals. The localisation of stable chromatin foci is depicted within the heat maps (top row, red pixels) and quantification of this population reveals the fraction of stable chromatin foci increase with time after DSB induction. (c) Quantification of the number of stable compact chromatin pixels across multiple cells (N=10, mean ± s.e.m.) in the first hour following NIR induced DSBs (dashed line represents average number of stable compact chromatin pixels in an unperturbed cell). (d) Chromatin compaction size map derived by PLICS analysis 0-30 min after DSB induction. (e) Histogram of size and distance changes induced by DSB in the data presented in (d). (f) Change in mean size and distance between compacted chromatin foci as a function of time during the DNA damage response (N=10 cells, mean ± s.e.m.). Scale bar equals 5 μm.

**Figure 5:**
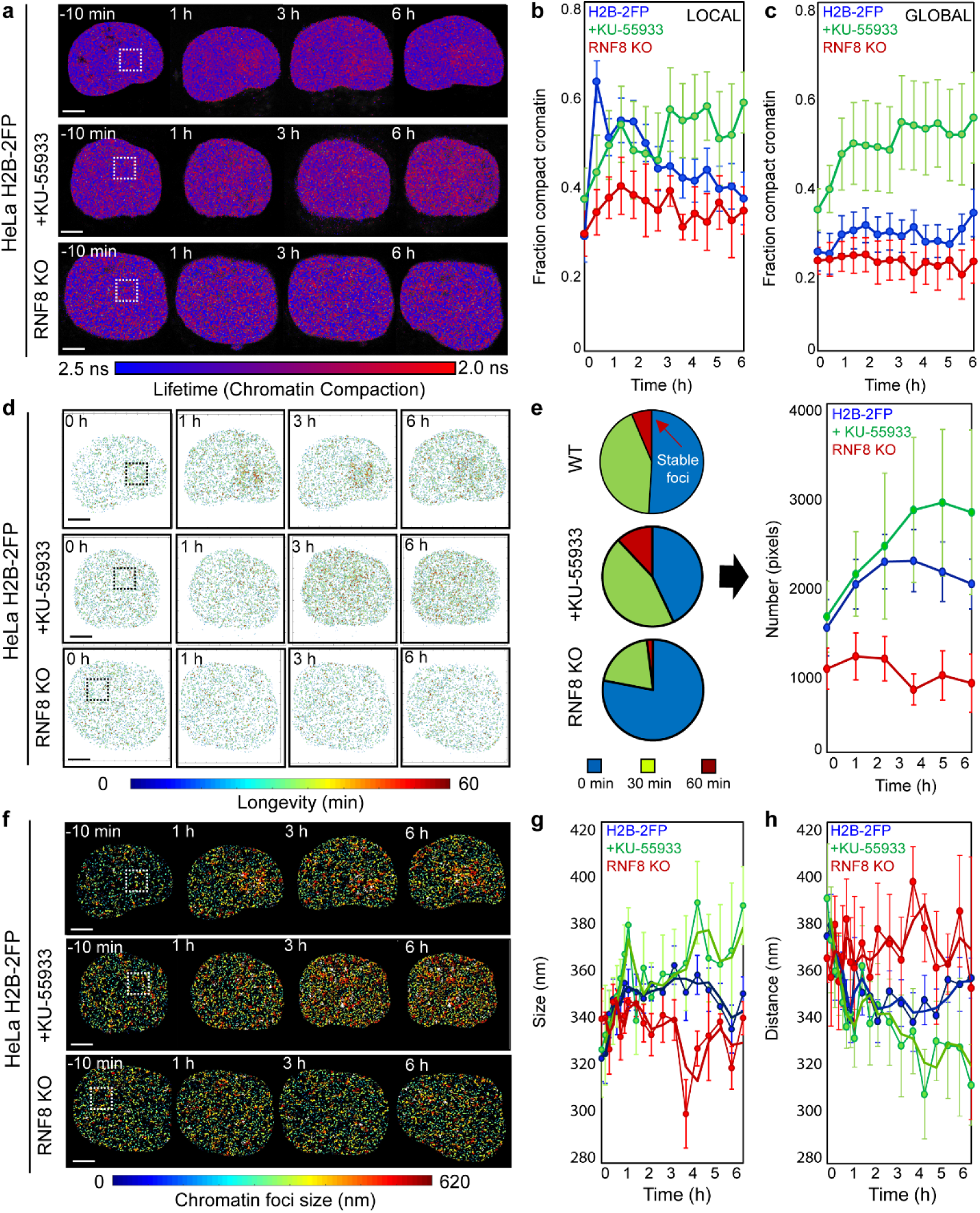
ATM and RNF8 regulate chromatin architecture in the DDR. (a) FLIM-FRET maps acquired in a HeLa^H2B-2FP^, HeLa^H2B-2FP^ cultures treated with KU-55933, or HeLa^H2B-2FP^ RNF8 KO cells, before and 1 h, 3 h and 6 h after micro-irradiation. (b, c) Fraction of compacted chromatin pixels within (b) and outside (c) of the DNA damage site in HeLa^H2B-2FP^ (blue curve, N=10), KU-55933 treated (green curve, N=6) or RNF8 KO cells (red curve, N=3) (mean ± s.e.m). (d) Longevity maps of compact chromatin foci from the cells in (a). (e) Quantification of compact chromatin foci stability (left) throughout during the DDR (right) in NIR laser treated HeLa^H2B-2FP^, KU-55933 treated HeLa^H2B-2FP^, and HeLa^H2B-2FP^ RNF8 KO cells (N as indicated above, mean ± s.e.m). (f) Chromatin compaction size map derived by PLICS from the cells shown in (a). (g, h) Change in the mean size (g), and distance between (h) of compacted chromatin foci during the DDR in NIR laser treated in HeLa^H2B-2FP^ (blue curve, N=10), KU-55933 treated (green curve, N=6) or RNF8 KO cells (red curve, N=3) (mean ± s.e.m). Scale bar equals 5 μm.

Applying PLICS and iPLICS to the same data set revealed that in the first 30 min following NIR laser irradiation there was an increase in the size of compact chromatin foci surrounding the NIR treated ROI (Fig. 4d-f). This local chromatin reorganisation gave rise to an overall increase in the detected compact chromatin foci size (left panel of Fig 4e) and a subtle reduction in the global spatial distribution of compacted chromatin foci during the first 30 min following DSB induction (right panel of Fig. 4e). Similar, to longevity analysis, PLICS and iPLICS demonstrate a uniform increase across multiple cells in compact chromatin foci size (320 nm up to 370 nm) and decrease in spacing between foci (505 nm down to 470 nm) within the first 30 min of the DDR (Fig. 4f). Thus, the stable chromatin border detected by longevity analysis at the perimeter of the DNA damage site arises from a local increase in chromatin foci size.

### ATM and RNF8 regulate DDR-dependent chromatin dynamics

To probe DDR-dependent regulation of chromatin architecture, we inhibited key DDR enzymes and measured the impact on chromatin dynamics following DSB induction. For these experiments we inhibited ATM kinase with the potent and specific inhibitor KU-55933 (54). Additionally, we used CRISPR/Cas9 to genetically delete RNF8 in HeLa^H2B-2FP^. ATM inhibition in HeLa^H2B-2FP^ cells was confirmed by western blot (Supplementary Fig. 6a) and *RNF8* deletion in HeLa^H2B-2FP^ RNF8 knockout (KO) cells was confirmed by sequencing the targeted locus and visualising suppressed 53BP1 recruitment to DSBs induced by ionizing or NIR laser irradiation (Supplementary Fig. 6b-e). We acquired phasor FLIM-FRET data in KU-55933 treated HeLa^H2B-2FP^ cultures and HeLa^H2B-2FP^ RNF8 KO cells, before and for six hours post NIR irradiation (Fig. 5a). ATM inhibition corresponded with an inhibition of acute chromatin compaction surrounding the lesion site within the first hour after break induction (green curve, Fig. 5b) and showed a gradual increase in global chromatin compaction throughout the experiment (green curve, Fig. 5c). *RNF8* deletion suppressed both local rearrangement of chromatin architecture at the lesion (red curve, Fig. 5b) and inhibited alteration of global chromatin compaction following break induction (red curve, Fig. 5c).

**Figure 6:**
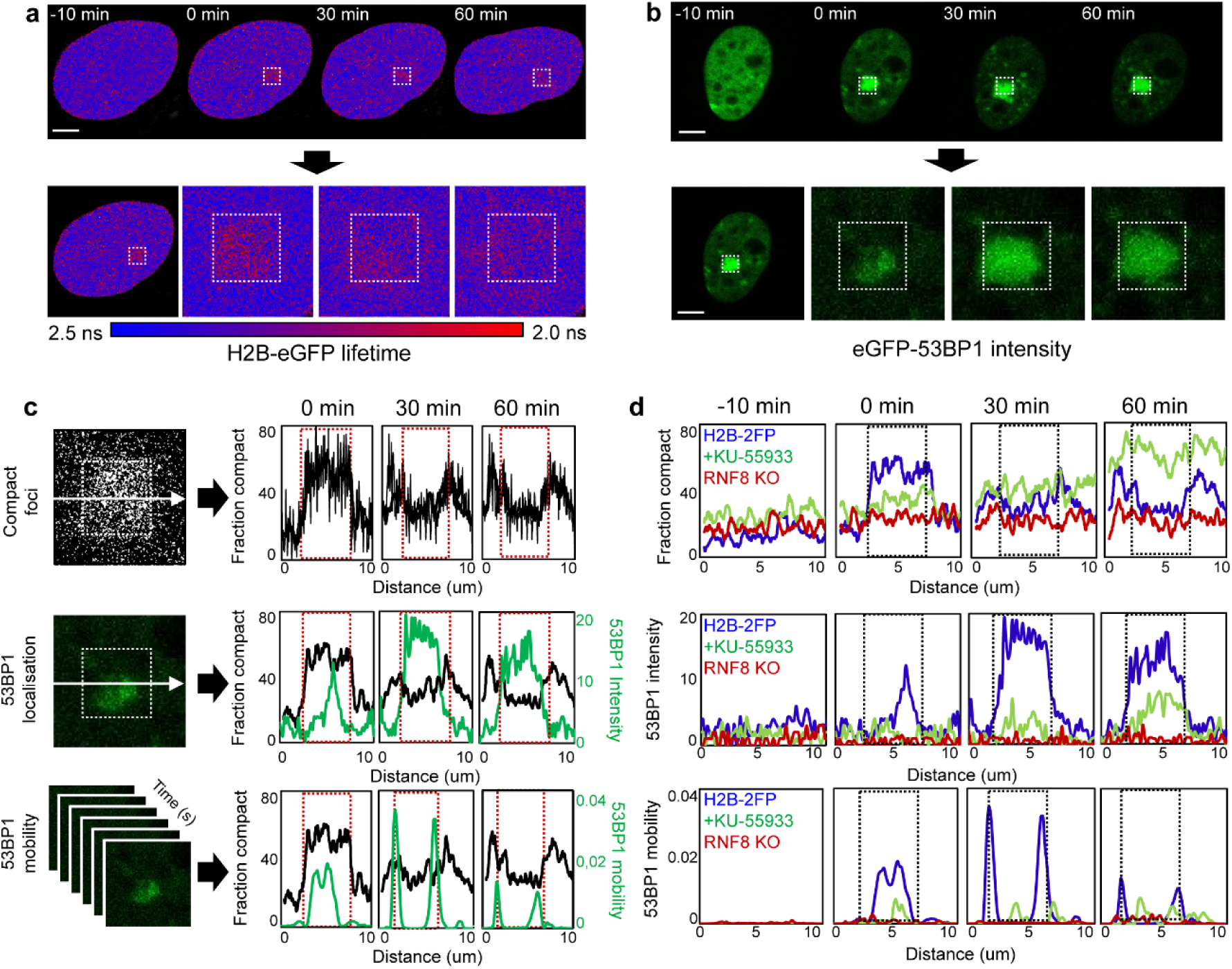
Chromatin architecture demarcates the repair locus. (a) FLIM-FRET maps acquired in a HeLa^H2B-2FP^ cells before and 0 min, 30 min and 60 min after micro-irradiation (top row) and expanded images of the DSB region of interest (bottom row) at these time points. (b) eGFP-53BP1 intensity images acquired in a HeLa cells before and 0 min, 30 min and 60 min after micro-irradiation (top row) and expanded images of the DSB region of interest (bottom row) at these time points. (c) Correlation of compact chromatin foci localisation along the horizontal axis as a function of time (top row, image from 0 min shown) with 53BP1 localisation (middle row, image from 0 min shown) and mobility (bottom row). eGFP-53BP1 localisation and mobility are also averaged along the horizontal axes (green plots). Red dashed box indicates laser micro-irradiation ROI. (d) Comparison of compact chromatin foci localisation (top row), eGFP-53BP1 localisation (middle row) and eGFP-53BP1 mobility (bottom row) in untreated HeLa cells (blue curve), KU-55933 treated HeLa cells (green curve) or RNF8 KO cells (red curve) before and 0 min, 30 min and 60 min after micro-irradiation. Representative example of N=3 shown.

Longevity analysis revealed that both ATM inhibition and *RNF8* deletion disrupted formation of the stable chromatin border surrounding the damage site (Fig 5d). ATM inhibition and RNF8 deletion also altered global compact chromatin stability over the six-hour imaging duration following DSB induction (Fig 5e). Specifically, KU-55933 conferred the gradual accumulation of stable compact chromatin foci throughout the nucleus with no geographical connection to the NIR treated ROI, while RNF8 deletion resulted in more dynamic chromatin throughout the experiment. Likewise, PLICS and iPLICS analysis revealed that both KU-55933 and *RNF8* KO suppressed the increase in compacted chromatin foci size in the first 30 min after break induction (Fig. 5f). Over the course of the experiment, ATM inhibition corresponded with a persistent and gradual increase in compact chromatin foci size and reduced spacing throughout the nucleus, irrespective of the damage site (green curves, Fig. 5g-h). Whereas *RNF8* KO cells fluctuated between significant increases and decreases in compact chromatin foci size and spacing, without exhibiting any trend (red curves, Fig. 5g-h). Together, these results demonstrate ATM and RNF8 to spatiotemporally regulate chromatin decompaction and compaction activities at the DSB locus and the surrounding global chromatin network organisation. We note that the nuclear wide gradual increase in chromatin compaction with KU-55933 treatment is a phenotype that after DSB induction continued independent of damage location. We anticipate this reflects an off-target effect of KU-55933, or a cellular response to acute ATM inhibition independent of the cellular response to DSBs.

### Chromatin architecture demarcates DSB repair locus boundaries

To determine how chromatin architecture at DSBs relates to the local DNA repair environment, we correlated the spatial localisation of compact chromatin in HeLa^H2B-2FP^ with eGFP-53BP1 mobility and localisation in parental HeLa cells. To do so, we measured 53BP1 mobility by pair correlation analysis: a branch of spatiotemporal correlation spectroscopy demonstrated previously in a study of Ku-GFP dynamics during DSB repair (35). Pair correlation is based on the acquisition of fluctuations in fluorescent intensity in each pixel of a line scan (34, 55, 56). From spatial cross correlation of pairs of fluorescent fluctuations located outside and inside a DSB site, the mobility of a fluorescently tagged DNA repair factor within or adjacent to a lesion can be determined (35). For these experiments, HeLa parental cells were transiently transfected with eGFP-53BP1. eGFP-53BP1 location and mobility were then monitored before and during the first hour after DSB induction via NIR laser micro-irradiation using the exact settings employed for the FLIM-FRET experiments above. The eGFP-53BP1 data in parental HeLa cells were then correlated to observed chromatin dynamics in HeLa^H2B-2FP^ (Fig. 6a, b). Quantitation of chromatin compaction along a line intersecting the NIR laser treated focus in HeLa^H2B-2FP^ confirmed that chromatin is rapidly compacted immediately after break induction, and that by thirty minutes the compact chromatin macrostructure had propagated outward to form the surrounding compact chromatin ring (Fig. 6c, top row). Immediately following break induction, eGFP-53BP1 accumulated at the NIR-laser treated locus, reaching maximum intensity by 30 min (Fig. 6c, middle row). Notably the radial spread of eGFP-53BP1 was constrained at the boundary of the surrounding compact chromatin border.

We then plotted 53BP1 movement by performing pair correlation across the same line intersecting the NIR laser treated focus. These data revealed increased eGFP-53BP1 mobility at the break immediately following DNA damage (Fig. 6c, bottom row). By thirty minutes, eGFP-53BP1 mobility was constrained to the interface of the compact chromatin border, with mobility declining by sixty minutes. Notably, eGFP-53BP1 mobility superimposes spatiotemporally with outward propagation of the compact chromatin border and spread of eGFP-53BP1 accumulation. Also, ATM inhibition and *RNF8* deletion in parental HeLa suppressed eGFP-53BP1 localisation and mobility at the damage site (Fig. 6d). Within the context of chromatin architecture, eGFP-53BP1 accumulated in the decompacted chromatin internal to the break while being excluded from the compact chromatin surrounding the laser treated ROI. We interpret these data to indicate that establishment of a transient compact chromatin border demarcates the repair locus from the surrounding chromatin environment.

## DISCUSSION

In this study we established a biophysical method to measure chromatin organisation in live cells using the phasor approach to FLIM-FRET analysis of fluorescently labelled histones. Coupling this technology with laser micro-irradiation and a novel analysis workflow enabled us to identify the DDR-dependent chromatin architectural changes that occur with DSBs. We found that ATM and RNF8 regulate chromatin decompaction at break sites, coupled with formation of a compact chromatin border that physically demarcates the repair locus and modulates 53BP1 mobility.

A major advance of this study is that the imaging and analysis tools we have developed enable local chromatin structure to be measured on a nuclear wide scale in a living cell, whilst maintaining sensitivity toward heterogeneity of individual chromatin foci dynamics during the DNA damage response. We achieved this through phasor analysis of FLIM and image correlation spectrums. Phasor provides a fit-free solution that does not require a priori knowledge of system complexity (40, 47). Instead, phasor analysis of histone FLIM-FRET directly maps the chromatin compaction status at each pixel, independent of the multitude of potential compaction states. This in turn enabled us to extract the localisation of specific chromatin foci, assess their temporal stability by longevity analysis, and characterise spatial organisation in terms of individual chromatin foci size or spacing by PLICS and iPLICS. Thus, by applying our technique to the DDR we could generate high-resolution spatiotemporal maps of compacted chromatin status surrounding DSBs which are not achievable through other analytical tools. While our method does not supply the sequence information that is provided by ChiP-seq or 3C-based methods, it does enable spatiotemporal chromatin compaction analysis in a single cell over time, which is not possible with current sequencing techniques.

Together our longevity and PLICS analysis revealed the uniform alteration of chromatin architecture within the first thirty minutes following damage induction, which is consistent with a highly regulated process. Resolution of chromatin architecture occurs thereafter with cell to cell heterogeneity over three to six hours. This is consistent with the kinetics of DDR focus resolution and in agreement with resolution of chromatin alteration at DSBs occurring with completion of repair. We anticipate resolution heterogeneity reflects differing cell to cell repair kinetics, dependent upon the genetic location where breaks occurred, the cell cycle, or other unknown factors (57-59). Critically, it suggests that DDR-dependent changes in chromatin architecture are tightly regulated through the repair process from break induction to repair completion. Additionally, these observations identify that our analysis is sufficiently sensitive to monitor single cell heterogeneity in chromatin macrostructure.

Correlation between our phasor FLIM-FRET and eGFP-53BP1 pair correlation analysis provided insight into the relationship between chromatin architecture and establishment of the repair focus. Immediately following break induction, 53BP1 is highly mobile at the damage site. By thirty minutes, 53BP1 accumulation had maximised and its mobility declined within the decompacted chromatin interior of the repair focus. At the same time, 53BP1 motility was observed only at the interface of the expanding compact chromatin border. These data are consistent with 53BP1 binding to the decompacted chromatin as this substrate becomes available with outward expansion of the compact chromatin border. Mobility of 53BP1 is high at the chromatin border as this protein continues to localise to the break, but its binding is restricted within the compact chromatin environment. We interpret these findings to signify that the compacted chromatin boundary serves to distinguish the decompacted repair locus from the remainder of the nucleus.

The significance of this physical separation of the repair locus from the surrounding environment may provide an important function to the DNA repair process. Chromatin decompaction and nucleosome eviction is proposed to ease access of DNA repair factors to their DNA substrate and increase repair kinetics (3, 20, 60). Conversely, the compacted chromatin border excludes 53BP1 retention, and may also inhibit the spurious binding of other DNA repair factors. We also predict the compacted chromatin border likely contributes to DDR-dependent transcriptional inhibition, which is predicted to halt passage of RNA polymerases through a DDR locus to enable efficient repair [14, 20]. DDR-dependent transcriptional silencing occurs in cis and is dependent upon ATM and RNF8 (11, 61). This is consistent with our observation that formation of the compact chromatin border surrounding the damage focus is ATM and RNF8-dependent.

Consistent with their key function in the DDR, we identified that ATM and RNF8 regulate chromatin architecture at NIR laser-induced DSBs. RNF8-dependent chromatin decompaction is also consistent with a role for this protein in regulating the local chromatin environment to promote assembly of checkpoint and repair machineries at DNA lesions (62). As RNF8 activity and chromatin ubiquitylation is essential for 53BP1 binding (18), our data suggest the interior decompacted chromatin environment coincides with the domain of ubiquitylated chromatin at the break. Cumulatively our data provide a macrostructural understanding of how chromatin architecture is arranged at the DDR focus. Further they imply that ATM and RNF8 regulation of a compact chromatin border serves to demarcate and arrange the repair locus to simultaneously balance DNA repair factor accumulation and potentially transcriptional shutdown of neighbouring genes.

In conclusion, phasor histone FLIM-FRET analysis coupled with laser micro-irradiation enabled us this study of DDR-dependent chromatin dynamics with unprecedent spatiotemporal resolution at the single cell level. Whilst we focused on the DDR in this study, the methodology presented here is agnostic to research topic, and as we have shown is easily combined with physical, pharmacological or genetic manipulation of the underlying system. As chromatin dynamics underlie a wide range of biological functions, we anticipate the analytical tools presented here are widely applicable and will contribute to advancing our understanding of how genomes work in the context of living cells.

## METHODS

### Plasmids for cell line and vector generation

pLXSN H2B-mCherry was as kind gift from Laure Crabbe (63). pWZL H2B-eGFP was created by PCR amplifying H2B-eGFP from eGFP-N1 H2B-eGFP (a gift from Geoff Wahl, Addgene plasmid #11680) (64) using the BamHI H2B-eGFP forward and EcoRI H2B-eGFP reverse oligonucleotides (Supplementary Table 1), followed by in-fusion (Clontech) cloning into BamHI and EcoRI (New England Biolabs) digested pWZL vector (a gift from Scott Lowe, Addgene #18750). pLentiCRISPRv2 was a gift from Feng Zhang (Addgene plasmid #52961) (65). pLentiCRISPRv2-RNF8-2 was generated by cloning an sgRNA sequence that targets exon 1 of RNF8 by annealing the sgRNF8-2 sense and antisense oligonucleotides (Supplementary table 1) and in-fusion cloning into BsmBI digested pLentiCRISPRv2. pcDNA5-FRT/TO-eGFP-53BP1was a gift from Dan Durocher (Addgene plasmid 60813) (18). All plasmids were verified by Sanger sequencing (Australian Genome Research Facility, Sydney).

### Cell Culture and treatment

HeLa cells were originally provided by Jan Karlseder (Salk Institute, La Jolla, CA). Cells lines were propagated in DMEM (Life Technologies) supplemented with 1% non-essential amino acids (Life Technologies), 1% glutamax (Life Technologies), and 10% bovine growth serum, at 37°C, 10% CO and, 3% O_2_. During FLIM-FRET imaging, cells were cultured in DMEM (Lonza), supplemented with 10% fetal bovine serum (Gibco), HEPES (Gibco) and 1x Pen-Strep (Lonza). Identity of all cell lines was confirmed using tandem repeat profiling by Cell Bank Australia (CMRI, Westmead), and cell lines were confirmed to be Mycoplasma negative (MycoAlert, LT07-118, Lonza). For experimentation to establish unquenched fluorescence lifetime of H2B-eGFP, identify laser micro-irradiation conditions for DSB induction, or measurement of DNA repair factor mobility, HeLa cells were transiently transfected with pWZL H2B-eGFP, Ku-GFP or eGFP-53BP1 using Lipofectamine 3000 according to manufactures instructions. ATM was inhibited with 10 µM KU-55933 (Merck Millipore). Chromatin compaction was induced with 5 μg ml^-1^ Actinomycin D, as shown previously to stop class III transcription (Sigma) (50). Chromatin decompaction was induced with 400 nM Trichostatin A (Abcam) (32). For RNF8 clone screening, cells were subjected to ionizing radiation using a Gammacell 3000 Elan (Best Theratronics).

### Generation of HeLa^H2B-2FP^

pLXSN H2B-mCherry and pWZL H2B-wGFP retrovectors were created by transfecting Phoenix-AMPHO cells (purchased from ATCC) with Lipofectamine 2000 according to manufactures instructions (ThermoFisher). Vector containing supernatant from pLXSN H2B-mCherry and pWZL H2B-eGFP co-transfected Phoenix-AMPHO cells were collected at 12 and 24 hours after transfection, passed through a 0.22 µm filter, and used to infect HeLa cells in the presence of 4 µg/ml polybrene (Sigma). Cultures were selected with 600 µg/mL G418 and 200 µg/mL Hygromycin before sorting at the Westmead Institute for Medical Research (WIMR) flow cytometry centre (Sydney, Australia). Cells with low expression of H2B-mCherry and H2B-eGFP were sorted, and propagated in 600 µg/mL G418, 200 µg/mL Hygromycin, and 1x Pen/Strep for 10 days before freezing multiple vials of cells for future experiments.

### Generation of RNF8 Knock out clones

pLentiCRISPRv2-RNF8-2 lentivectors were created in the Children’s Medical Research Institute Vector and Genome Engineering Facility (Sydney, Australia) as described previously (66). RNF8 was deleted in HeLa or HeLa^H2B-2FP^ by transducing cultures with pLentiCRISPRv2-RNF8-2 vectors at a MOI of 100 and selecting with 1 µg/mL puromycin for two days. Single cells were plated into individual wells in 96-well plates at the WIMR Flow cytometry centre, and cultures expanded for 14 days before freezing. Following continued expansion, clones were screened for inhibited 53BP1 recruitment to γ-H2AX foci by irradiating cells with 1 GY ionizing radiation and immunofluorescence staining as described below. Clones negative for 53BP1 recruitment were screened for genetic alteration by PCR amplifying the targeted locus using the RNF8 screen infusion forward and reverse primers and cloning the PCR products into PmeI (New England Biolabs) digested pcDNA3.1 (Thermofisher) using Infusion cloning. Plasmids were sequenced with RNF8 sequence forward, and Clones C4 and D5 verified to contain gene disruptive mutations in all three RNF8 alleles in HeLa were used for further experimentation.

### FLIM-FRET microscopy

All FLIM-FRET data was acquired with a Zeiss LSM 880 laser scanning microscope coupled to a 2-photon Ti:Sapphire laser (Spectra-Physics Mai Tai, Newport Beach) producing 80 fs pulses at a repetition of 80 MHz and a Picoquant Picoharp (Picoquant, Germany) for time resolved detection. A 60 X water immersion objective 1.2 NA (Zeiss, Germany) was used for all experiments. The 2-photon excitation laser was tuned to 900 nm for selective excitation of the donor fluorophore and minimisation of DNA damage during live cell imaging. A short pass 760 nm dichroic mirror was used to separate the fluorescence signal from the laser light. The fluorescence signal was directed through a 550 nm long pass eGFP / mCherry filter, and the donor and acceptor signal split between two photomultiplier detectors (H7422P-40 of Hamamatsu), with the following bandwidth filters placed in front of each: eGFP 520/25 and mCherry 590/25, respectively. We simultaneously acquire intensity and lifetime data by raster scanning the two-photon Ti:Sapphire laser beam (900 nm) across a selected HeLa^H2B-2FP^ interphase nucleus at a digital zoom that results in a frame size of 30 μm. The pixel frame size was set to 512, which gave a pixel size of 0.06 μm.

The scanning speed was set at 32.77 µs/pixel, which resulted in a frame time of 1.9 s. For each FLIM experiment 40-50 frames were integrated. Together these conditions resulted in an acquisition time of ∼2 min. Calibration of the system and phasor plot space was performed by measuring fluorescein (pH 9.0), which has a known single exponential lifetime of 4.04 ns. The FLIM data were acquired and processed by the SimFCS software developed at the Laboratory for Fluorescence Dynamics (LFD) (www.lfd.uci.edu).

### FLIM-FRET analysis

The fluorescence decay recorded in each pixel of a FLIM image was quantified by the phasor approach to lifetime analysis. As described in previously published papers, the decay in each pixel of a FLIM frame image is transformed into its sine and cosine components which are then represented in a two-dimensional polar plot (phasor plot), according to equations 1 and 2:

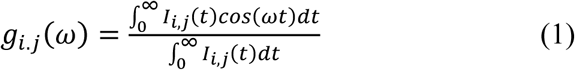

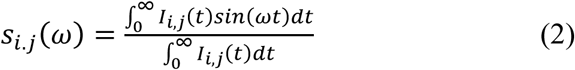

where ω is the laser repetition angular frequency (2πf) and the indexes *i* and *j* identify a pixel of the image. Each pixel of the FLIM image thus gives rise to a single point (phasor) in the phasor plot and when used in reciprocal mode, enables each point of the phasor plot to be mapped to each pixel of the FLIM image. Since phasors follow simple vector algebra, it is possible to determine the fractional contribution of two or more independent molecular species coexisting in the same pixel. Thus, in the case of two independent species, all possible weightings give a phasor distribution along a linear trajectory that joins the phasors of the individual species in pure form.

In the case of a FRET experiment where the lifetime of the donor molecule is changed upon interaction with an acceptor molecule, the realisation of all possible phasors quenched with different efficiencies describe a curved trajectory in the phasor plot. The experimental position of the phasor of a given pixel along the trajectory determines the amount of quenching and therefore the FRET efficiency of that location. The contributions of the background and of the donor without acceptor are evaluated using the rule of the linear combination, with the background phasor and the donor unquenched determined independently. In the case of HeLa^H2B-2FP^ the FRET efficiency varies as a function of local nucleosome proximity, thus the trajectory of phasor clusters between the donor phasor and FRET state of the biosensor represent the varying contributions of donor fluorescence and quenched donor fluorescence in any one pixel. By moving the phasor cursor between these two terminal phasor locations, we can calculate the exact FRET efficiency and therefore local chromatin compaction status for each pixel highlighted. The FLIM-FRET analysis was performed in the SimFCS software developed at the LFD (www.lfd.uci.edu).

### Temporal longevity analysis

A phasor FRET map reports the localisation of a range of chromatin compaction states detected within a FLIM image. Thus, by extracting the x-y coordinates of pixels only in a low or high FRET state we can derive a binary image of ‘open’ or ‘compacted’ chromatin localisation. If we are interested in the temporal persistence of each located chromatin foci, we can generate a time series of chromatin localisation maps and from application of a moving average (MAV) across them, derive a heat map of pixel longevity for the selected MAV temporal window. The number of frames employed for the MAV determines the range of amplitudes recorded in a longevity map and thus the threshold parameters for extracting the localisation of stable versus transient chromatin foci. For example, in Fig. 3 we average 3 frames acquired with a 10 min interval and the resulting longevity map contains pixels with an amplitude of 0, 0.333, 0.666 or 1. These values correspond to chromatin foci that are either not present in a pixel or present for 1, 2 or 3 frames, which in turns corresponds to chromatin foci that persist from 0 up to 20 min. By extracting the x-y coordinates of pixels only displaying an amplitude of 0.333 or 1 we can derive a binary image of transient versus stable chromatin foci for further spatial analysis. By counting the number of pixels present for 0-3 frames we can obtain a histogram which reports overall chromatin network stability. Depending on the acquisition procedure employed for generation of a time series of phasor FRET maps, we can also tune the analysis to extract fast or slow timescale dynamics. For example, in Fig. 4-5 we demonstrate that if we employ a MAV of three frames acquired with a 10 min interval this is suitable to interrogate chromatin border formation local to the DNA damage site. But if we instead employ a MAV of three frames across 1 h (0, 30 and 60 min) this temporal window is sensitive to changes in global chromatin organisation upon DSB enzyme inhibition and during the entire 6 h DDR. The chromatin localisation maps were generated from SimFCS and the longevity analysis routine, which can extract when, where and how many chromatin foci are stable within a time series of chromatin localisation maps, was implemented in Matlab (The Mathworks, Natick, MA).

### PLICS analysis

To investigate the size and spacing between chromatin foci localised in a phasor FRET map, similar to longevity analysis, we first extracted the x-y coordinates of pixels in an ‘open’ or ‘compacted’ chromatin state. Then from the resulting binary images we applied the PLICS approach. PLICS (Phasor analysis of Local Image Correlation Spectroscopy) consists of computing the spatial autocorrelation function (ACF) of a small region of the image, compute its angular average and transform the resulting curve in the phasor space. Successively, the phase coordinate of the transformed local ACF is stored and is used to characterize the ACF shape, which is linked to the size of the local structures. This approach is iteratively applied in order to analyse the entire image, providing a mapping of the size of the structures. Since the analysis involves binary images, the distance between two structures is merely the amount of empty space between them, therefore applying PLICS to the negative of the binary image provides a direct measurement of the distance between the structures. We will refer to this analysis as to iPLICS (inverse PLICS). The average values are displayed as the mean and standard deviation of the values of the size and distance maps. For the binary images analysed in this work, a 12 × 12 PLICS moving mask is used in order to obtain local size of the binary regions, while the negative of the binary image has been analysed with a 36 × 36 moving mask in order to characterize the distance between the structures. A dedicated calibration has been employed to extract the size of the binary regions by simulating binary circles with varying diameter, the details of the calibration procedure for an arbitrary function and the complete characterization of the technique are described in (47). The chromatin localisation maps were generated from SimFCS and the PLICS / iPLICS analysis routine, which can size and spacing of chromatin foci within a time series of chromatin localisation maps, was implemented in Matlab (The Mathworks, Natick, MA).

### Laser micro-irradiation and FLIM-FRET microscopy during the DDR

For all micro-irradiation experiments the 2-photon Ti:Sapphire laser (80fs, repetition rate 80MHz, Spectra-Physics Mai Tai, Newport Beach) was tuned to 780nm and used in conjunction with the Zeiss LSM880 laser scanning microscope (51). The laser beam was then focused on a small section of the nucleus (3 µm × 3 µm) which avoided the nucleolus or nuclear envelope and a frame scan acquired (256 × 256 pixels, 32.77µs/pixel, total acquisition time 1.15s) at a power found to recruit DNA repair factor Ku-GFP or eGFP-53BP1 (Supplementary Fig. 2). FLIM-FRET microscopy was performed in parallel using the microscope and acquisition settings described above. FLIM-FRET acquisitions were recorded before laser micro-irradiation at the following time points after this treatment: (1) 10 minute interval during the first hour of the DDR (0, 10, 20, 30, 40, 50, 60 min), (2) 20 minute interval during the second hour of the DDR (80, 100, 120 min) and (3) 30 minute interval for the remaining 4 hours of the DDR (150, 180, 210, 240, 270, 300, 330 and 360 minutes).

### Laser micro-irradiation and live cell imaging of eGFP-53BP1 mobility during the DDR

The Zeiss LSM880 laser scanning microscope was employed for live cell imaging of eGFP-53BP1 localisation and mobility. Specifically, a 60 X water immersion objective 1.2 NA (Zeiss, Germany) was used for all experiments and then a time series of eGFP-53BP1 intensity was acquired using the following microscope settings: 64 × 64 pixels, 8.19 μs/pixel, 2048 frames. eGFP was excited with the 488 nm emission line of an Argon laser and detected using the 510-560 nm emission range. The pair correlation analysis was performed in the SimFCS software developed at the LFD (www.lfd.uci.edu).

### Wide-field live cell imaging of HeLa^H2B-2FP^

Live cell imaging shown in Supplementary figure 1 was performed on a ZEISS Cell Observer inverted wide field microscope with a ZEISS HXP 120C mercury short-arc lamp and compatible filter cubes, using a 20x 0.8 NA air objective. Cells were plated on glass-bottom MatTek 6-well plates and images captured every six minutes for sixty-five hours at 37°C, 10% CO_2_, and atmospheric oxygen. Mitotic duration and outcome were scored by eye using Zen software (ZEISS).

### Immunofluorescence

Cells cultured on Alcain blue-stained glass coverslips were fixed with 2% paraformaldehyde in 1x PBS at room temperature for 10 min. The cells were rinsed in 1x PBS, the permeabilized in KCM buffer for 10 minutes (120 mM KCl, 20 mM NaCl, 10 mM Tris pH 7.5, 0.1% Triton X-100). Samples were blocked and RNAse treated in ABDIL (20 mM Tris pH 7.5, 2% BSA, 0.2% Fish Gelatin, 150 mM NaCl, 0.1% Triton, 0.1% Sodium Azide) containing 100 µg/mL RNAseA (Sigma) plus 2% normal goat serum (v/v) for one hour at room temperature. Samples were overlayed with 1:1000 primary antibody in ABDIL overnight at 4°C in a humidity chamber. The following day the cells were washed three times for five minutes with shaking in 1xPBS+1% tween (1x PBST), before overlaying with 1:1000 secondary antibody in ABDIL plus 2% normal goat serum (v/v) for one hour at room temperature. The slides were washed again as described above with DAPI added to the second wash at 50 ng/ml. Slides were dehydrated through a 70, 90 and 100% ethanol series by washing for 3 min in each condition, then air dried and mounded with prolong gold (Thermofisher). Images were captured on a ZEISS AxioImager Z.2 with a 63x 1.4 NA oil objective, appropriate filter cubes and an Axiocam 506 monochromatic camera using Zen software (ZEISS).

### Western Blots

Cells were collected with trypsin and the reaction quenched with growth media containing serum. Cells were washed in 1x PBS and homogenized in NuPage 4x LDS sample buffer without EDTA (Invitrogen) supplemented with 2% (v/v) β-mercaptoethanol (Sigma) and 5% (v/v) Benzonase (Merck Millipore) at 1 × 10^4^ cells/µL for 1 hr at room temperature. Cell lysate was denatured at 68°C for 10 minutes prior to resolution at 10 µL/well on Nu-PAGE 4-12% Bis-Tris gradient gels according to manufacturer’s protocols. Protein was transferred to nitrocellulose (Amersham) at 100 V for 1 hr and the membranes blocked for 1 hr with 5% (w/v) powdered skim milk in 1x PBS + 0.1% (v/v) Tween 20 (1x PBST). Membranes were probed overnight with primary antibody diluted in 1x PBST + 5% (w/v) powdered skim milk at 4°C with gentle nutation. The following morning blots were washed three times for 10 minutes with 1x PBST then probed with Horse Radish Peroxidase-conjugated secondary antibody diluted in 1x PBST + 5% (w/v) powdered skim milk for one hour at room temperature, followed by three washes in 1x PBST. Membranes were rinsed in deionized water blotted dry and overlaid with 0.22 µm filtered Enhanced Chemiluminescent western blotting substrate for 5 minutes (Pierce) before visualization with a Fujifilm LAS 4000 luminescent image analyser and associated software (GE Health).

### Oligonucleotides

Please see supplementary table 1 for a list of the oligonucleotides used in this study. All oligonucleotides were produced by Integrated DNA Technologies (Singapore).

### Antibodies

The following antibodies were used in this study: ATM (Cell Signalling Technology, 2873), ATM-S1981 (Abcam, ab81292), CHK2 (Millipore, 05-649), CHK2-T68 (Cell Signalling Technology, 2661), Vinculin (Sigma, V9131), γ-H2AX (Millipore, 05-636), 53BP1 (Santa Cruz, Sc-22760), Goat-anti-mouse Alexa488 conjugate (ThermoFisher, A11031), Goat-anti-rabbit Alexa 568 conjugate (ThermoFisher, A-11011), Polyclonal Goat Anti-Rabbit Immunoglobulins HRP-conjugate (Dako, P0448), Polyclonal Goat Anti-Mouse Immunoglobulins HRP-conjugated (Dako, P0447).

### Statistics and Figure preparation

Statistical analysis shown in Figure 4-6 and Supplementary Figure 1, 4 was performed using Graph Pad Prism (GraphPad Software). Sanger sequencing traces were exported from SnapGene. Figures were prepared using Adobe Photoshop, Illustrator, MatLab and SimFCS.

## ACKNOWLEDGEMENTS

All data are archived at University of Melbourne or the Children’s Medical Research Institute (CMRI). We thank the Biomedical Imaging Facility and the Mark Wainwright Analytical Centre at the University of New South Wales for enabling access to the Zeiss LSM 880. We thank Scott Page and the CMRI ACRF Telomere Analysis Centre supported by the Australian Cancer Research Foundation for assistance with imaging infrastructure. Leszek Lisowski and the CMRI Vector and Genome Engineering Facility are thanked for making Lentivectors. We thank the Westmead Institute for Medical Research Flow Cytometry Centre supported by the Cancer Council NSW and the Australian National Health and Medical Research Council (NHMRC) for cell sorting. K.G. is supported by grants from the Australian NHMRC (APP1059278) and Australian Research Council (ARC) (CE140100011 and LP140100967); E.G. is supported by grants from the US National Institute of Health (P41-GM103540 and P50-GM076516); A.J.C. is supported by grants from the Australian NHMRC (1053195, 1106241, 1104461), the Cancer Council NSW (RG 15-12), and the Cancer Institute NSW (11/FRL/5-02); and EH is supported by grants from the Australian NHMRC (1104461 and 1124762), Cancer Council NSW and ARC.

## SUPPLEMENTARY ITEMS

Jieqiong Lou^1^*, Lorenzo Scipioni^2^*, Belinda K. Wright^3^, Tara K. Bartolec^4^, Jessie Zhang^4^, V. Pragathi Masamsetti^4^, Katharina Gaus^3,5^, Enrico Gratton^2^, Anthony J. Cesare^4,†^, Elizabeth Hinde1,3,5,†.

Supplementary Figures 1-7

Supplementary Table 1

**Supplementary Figure 1:**
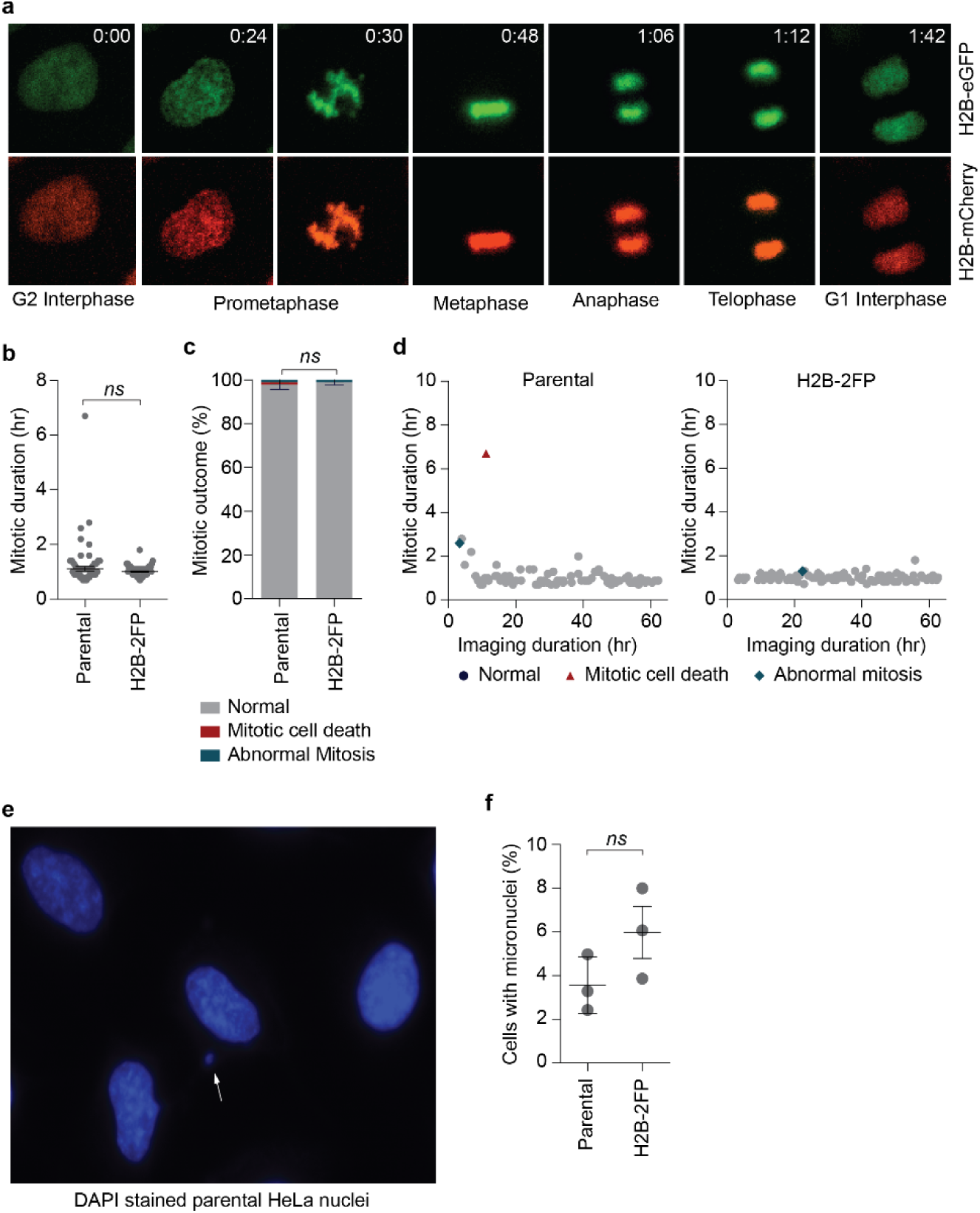
HeLa^H2B-2FP^ verification and assessment of genome stability. a) Incorporation of H2B-eGFP and H2B-mCherry into chromatin was verified by live cell imaging of HeLa^H2B-2FP^passing through mitosis. b-d) Mitotic phenotypes were measured in HeLa parental vs HeLa^H2B-2FP^ cells as a measure of genome stability. Mitotic duration (b) and outcome (c) were not impacted by exogenous histone expression in HeLa^H2B-2FP^ [*n =* 2 biological replicates of ≥ 40 mitoses per replicate, grouped into a single Tukey Box plot (b), or shown as a histogram mean ± range (c). *ns* = not significant, Mann-Whitney Text (b), Fisher Exact Test (c)]. d) Two-dimensional plots of the data from (b, c). Each point represents a mitotic event. Location of a symbol relative to the x-axis indicates time that mitosis started. Height on the y-axis represents the mitotic duration of that mitosis. The symbol identifies mitotic outcome. e) Image of DAPI stained HeLa parental cells with a micronucleus identified by an arrow. f) Quantitation of the percentage of cells containing micronuclei in HeLa parental and HeLa^H2B-2FP^ cultures (mean ± s.d., *n =* 3 biological replicates quantifying ≥ 445 nuclei per replicate, *ns* = not significant, student t-test).

**Supplementary Figure 2:**
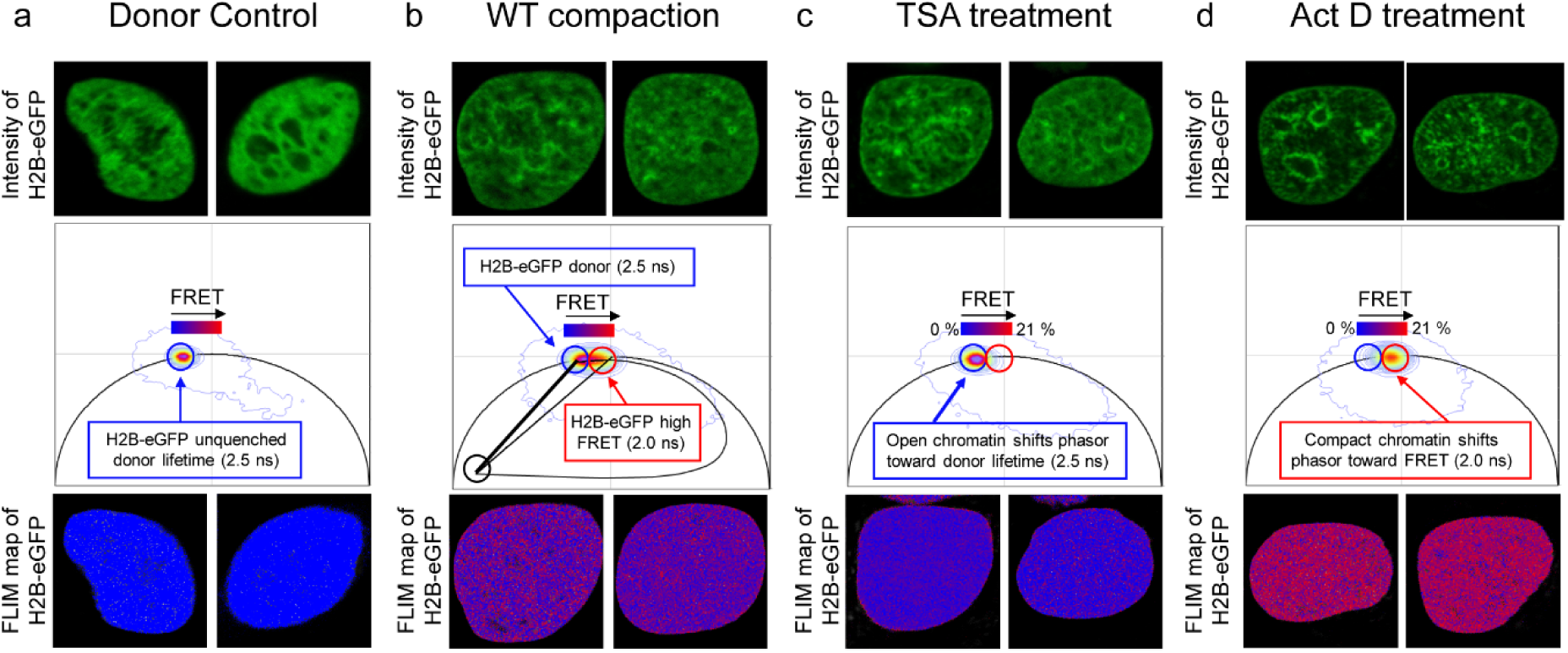
Establishment of FRET efficiency and phasor location of open and compacted chromatin in HeLa^H2B-2FP^. (a) FLIM-FRET analysis of a HeLa nucleus expressing H2B-eGFP in the absence of H2B-mCherry to establish the unquenched fluorescence lifetime and phasor location of H2B-eGFP (0% FRET, starting position of FRET trajectory) (τ_H2B-eGFP unquenched_ = 2.5 ns). (b) FLIM-FRET analysis of H2B-eGFP in a HeLa^H^2^B-2FP^ to establish the dynamic range of FRET efficiencies detectable in our biological system (0-21% corresponds to τ_H2B-eGFP_ = 2.0 - 2.5 ns). (c) FLIM-FRET analysis of H2B-eGFP in a HeLa^H2B-2FP^ treated with Trichostatin A (TSA) to identify the phasor location of open chromatin (0% FRET, τ_H2B-eGFP open chromatin_ = 2.5 ns). (d) FLIM-FRET analysis of H2B-eGFP in a HeLa^H2B-2FP^ treated with Actinomycin D (Act D) to identify the phasor location of compact chromatin (21% FRET, τ_H2B-eGFP compact chromatin_ = 2.0 ns).

**Supplementary Figure 3:**
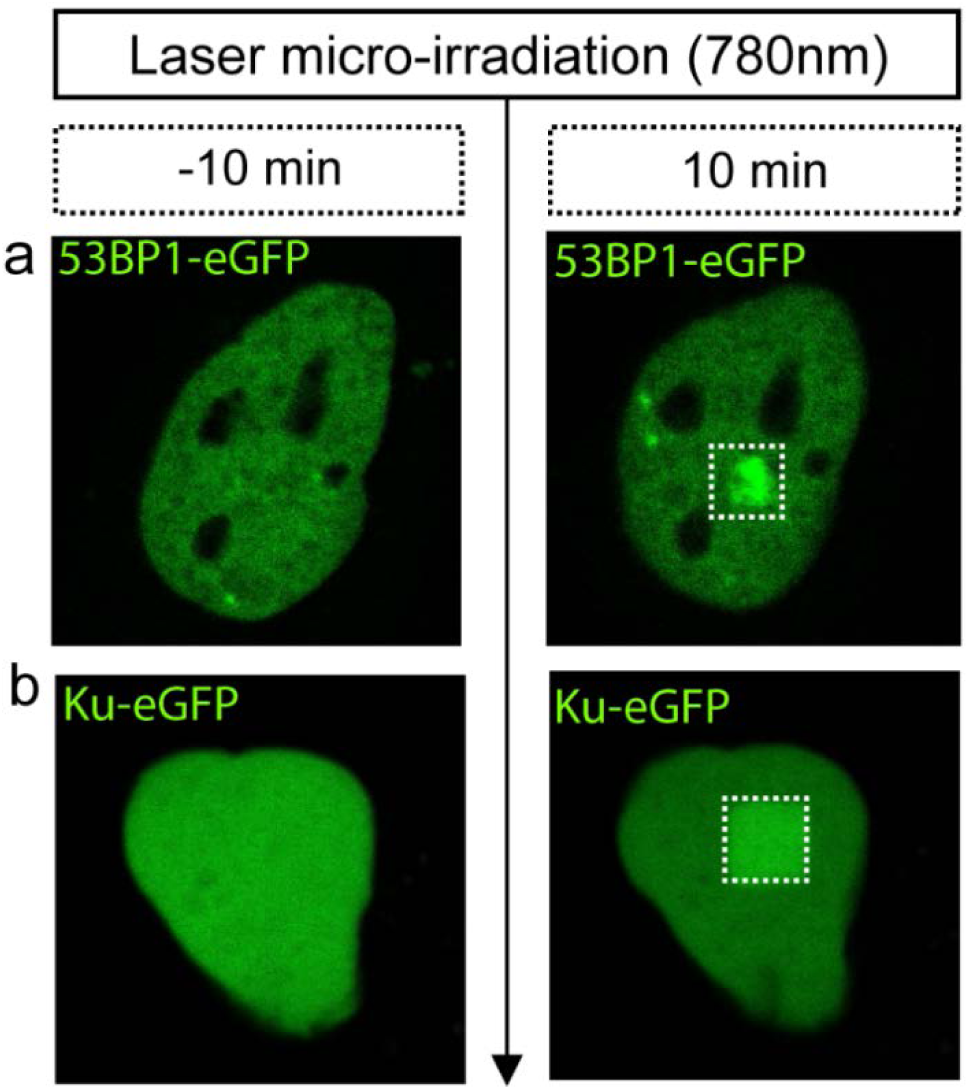
NIR laser micro-irradiation conditions optimized for double strand break (DSB) induction. Laser micro-irradiation of a HeLa nucleus expressing 53BP1-eGFP or Ku70-GFP with the Ti Sapphire 2-Photon laser operated at 780nm, 0.5mW for 1.15s results in damage site recruitment after 10 min.

**Supplementary Figure 4:**
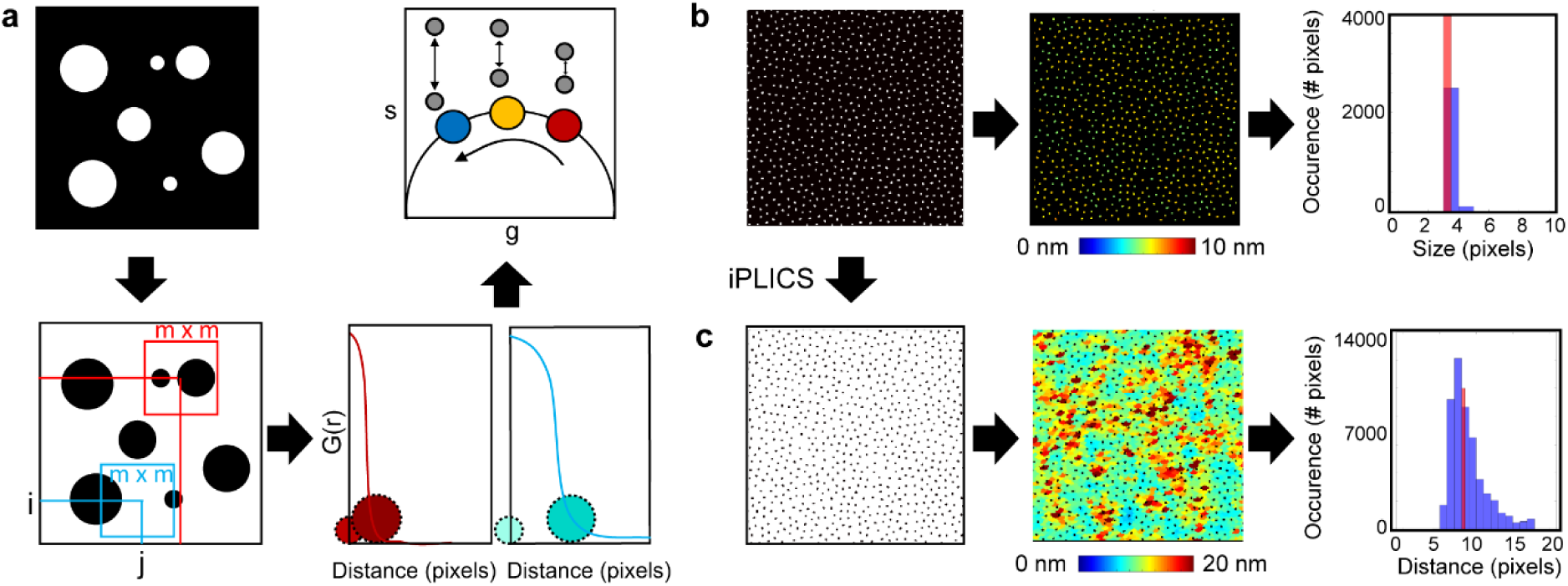
iPLICS analysis and a simulation to demonstrate the feasibility of quantifying chromatin foci size and spacing. (a) Schematic of iPLICS analysis: iPLICS starts with taking the negative of a chromatin localisation map, and then analogous to PLICS analysis, calculates localised two-dimensional spatial correlation functions that when collapsed into one-dimensional correlation profiles, describe the characteristic spacing between structures within each m × m matrix. Accordingly, by transforming each decay into phasor coordinates (g, s), we can graphically pseudo-colour each pixel according to spacing. (b) Simulation of circular binary structures with a diameter of 3 pixels and average distance between them of 8 pixels (left panel) enables derivation of a size map (middle panel) and the histogram of sizes that are pseudo-coloured (right panel). (c) Negative image of the simulation described in (b) enables inverse PLICS (iPLICS) analysis and therefore derivation of a distance map (middle panel) and the histogram of distances that are pseudo-coloured (right panel).

**Supplementary Figure 5:**
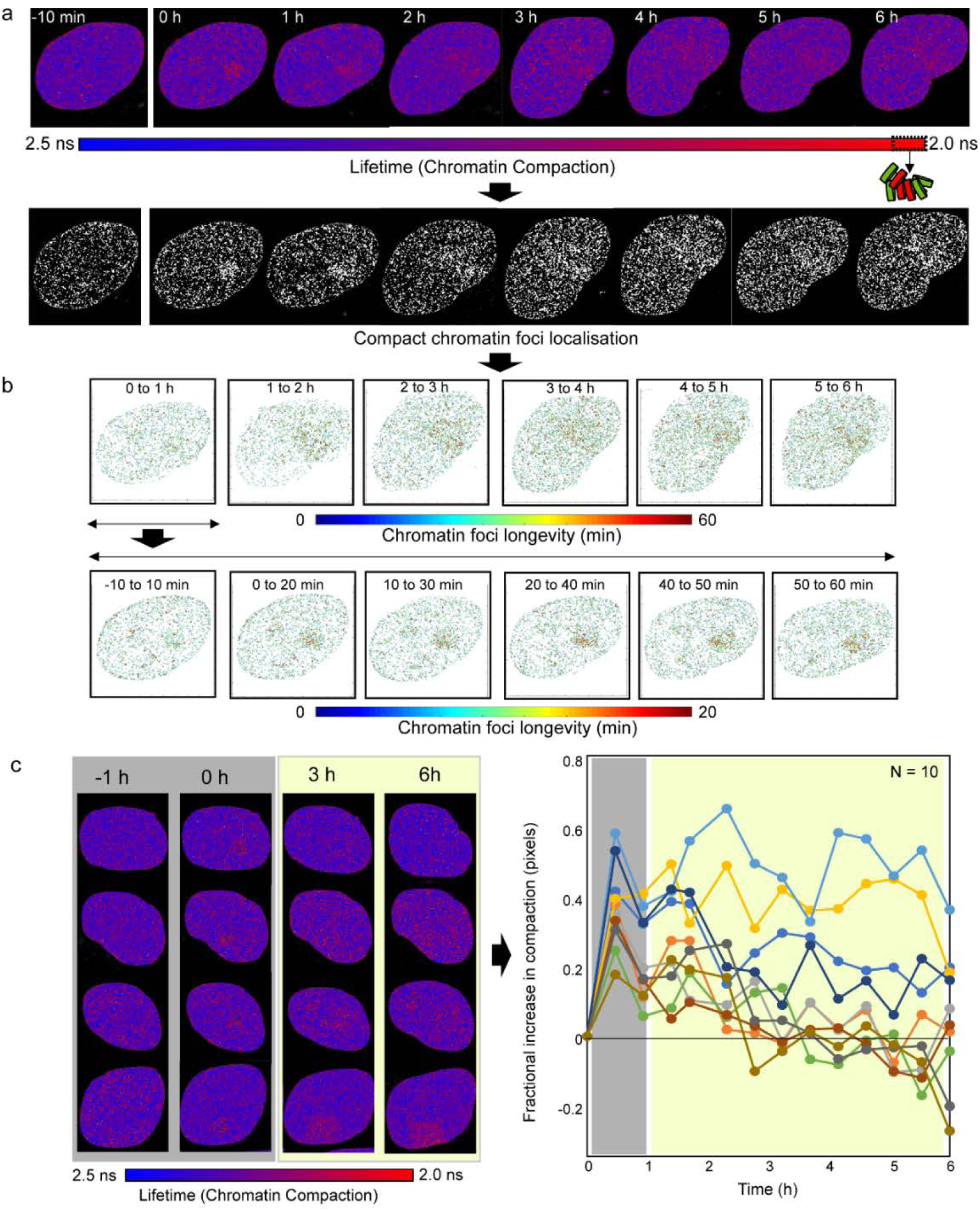
Heterogeneous resolution of the chromatin architecture after NIR irradiation. (a) FLIM-FRET maps acquired in a HeLa^H2B-2FP^ cell before and for 6 h after NIR irradiation (above) and their corresponding derived compact chromatin maps (below) localisation maps. (b) Longevity analysis of the full time series of compact chromatin localisation maps from (a) centred a ever hours (top row) or every ten minutes for the first hour (bottom row). (c) Left: Phasor maps from four examples of HeLa^H2B-2FP^ nuclei following NIR laser irradiation (four examples from N =10 cells). Right: quantitation of the fractional increase in compact chromatin across 10 HeLa^H2B-2FP^ cells following NIR laser irradiation. The grey boxes identify homogenous establishment of chromatin architecture following DSB induction, and the green shaded box the heterogenous resolution. Scale bar equals 5 μm.

**Supplementary Figure 6:**
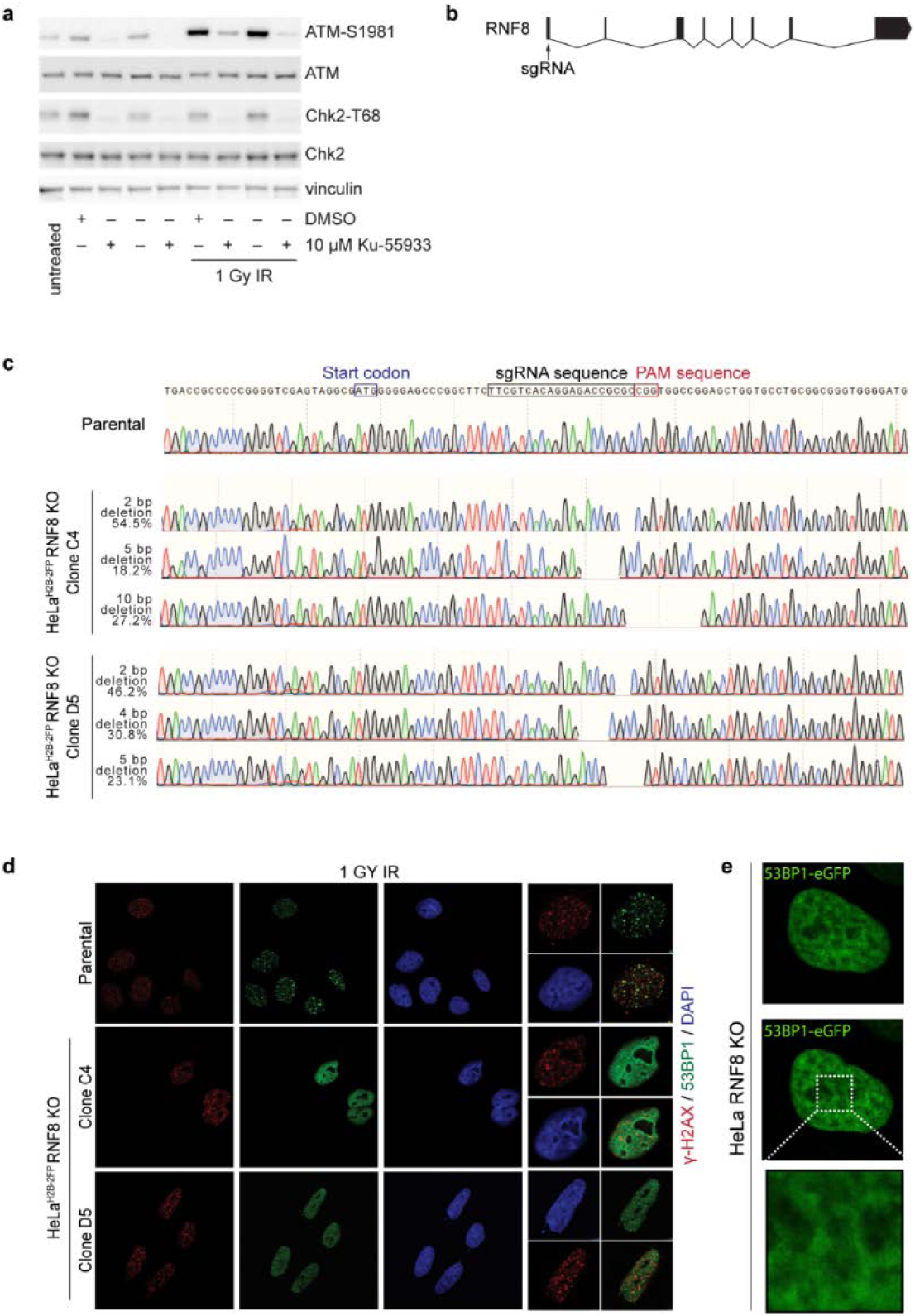
Verification of ATM inhibition and RNF8 knockout HeLa^H2B-2FP^. a) Western blots of whole cell extracts from HeLa^H2B-2FP^ cultures treated with vehicle (DMSO) or KU-55933, ± 1 Gray ionizing radiation (IR). 70,000 Cell equivalents were loaded per lane. b) Gene architecture of RNF8 and targeting of exon 1 by CRISPR/Cas9. c) Sequencing data of HeLa parental and CRISPR/Cas9 generated HeLa^H2B-2FP^ RNF8 KO clones. The start codon, targeting sgRNA, and PAM sequence are shown. HeLa cells contain three alleles of RNF8. HeLa^H2B-2FP^ RNF8 KO Clone C4 contained frame shifts inducing deletions of 2, 5 or 10 basepairs, while Clone D5 contained frameshift inducing deletions of 2, 4 and 5 basepairs. In order to resolve the genotype of mixed indel populations, a region surrounding the sgRNA targeted site was PCR amplified, subcloned into pcDNA3.1 and transformed into E.coli. 11 and 9 individual clones were sequenced for Clone C4 and D5 respectively. The percentage of clones with each targeted sequence are shown to the left of the sequencing traces. d) Immunofluorescence verification of RNF8 knockout. RNF8 is essential for 53BP1 recruitment to γ-H2AX-labelled double strand breaks induced by ionizing radiation. HeLa^H2B-2FP^, and HeLa^H2B-2FP^ RNF8 KO Clones C4 and D5 were treated with 1 GY IR, fixed and stained for γ-H2AX and 53BP1. 53BP1 localization to γ-H2AX foci is evident in HeLa^H2B-2FP^ but not the RNF8 KO clones. H2B-mCherry and H2B-eGFP are expressed at low levels enabling visualization of the strong γ-H2AX and 53BP1 IF staining following irradiation. e) Laser micro-irradiation of a HeLa RNF8 KO nucleus expressing 53BP1-eGFP under the same conditions identified in Supplementary Figure 2.

**Supplementary Figure 7:**
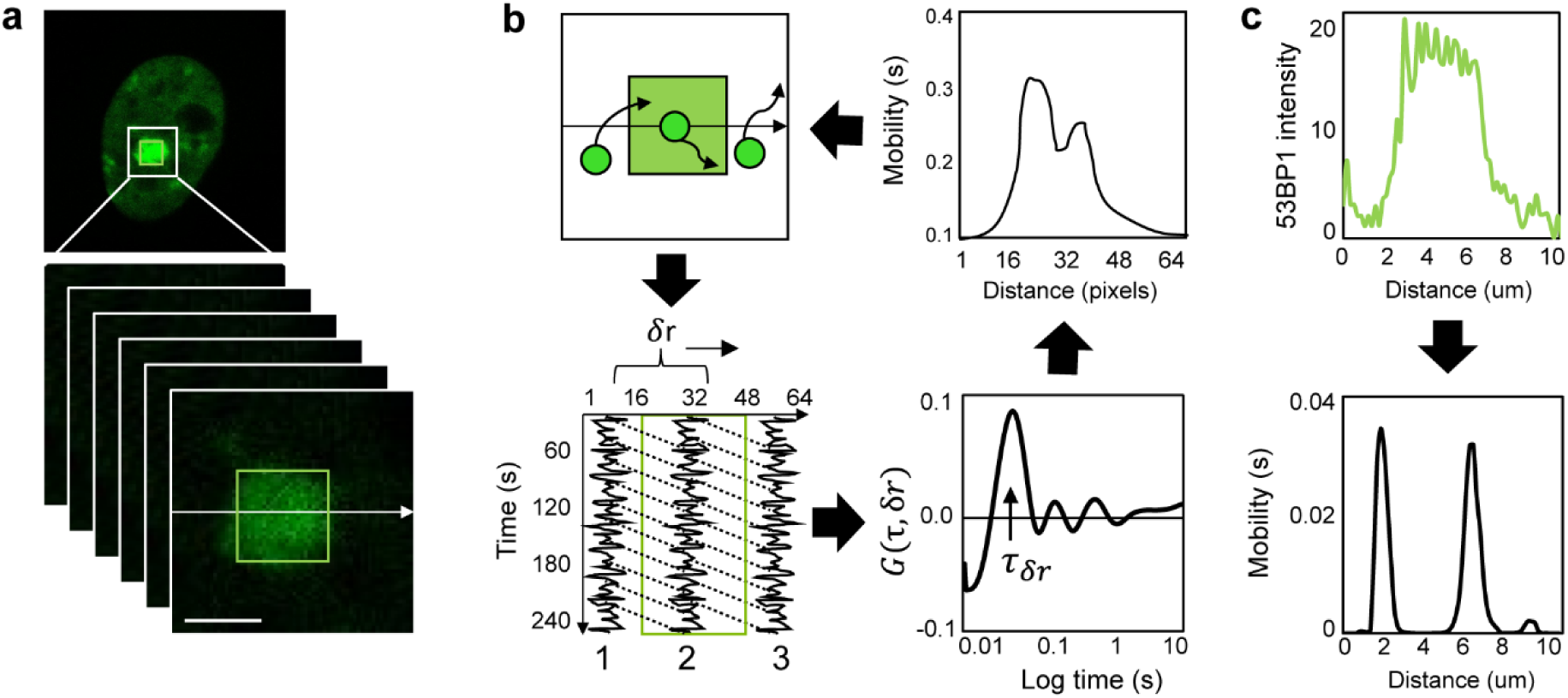
Pair correlation analysis of eGFP-53BP1 mobility. (a) eGFP-53BP1 intensity image acquired in a HeLa cell 30 min after micro-irradiation (top panel) and the region of interest (white box) sampled as a function of time for pair correlation analysis of eGFP-53BP1 mobility (bottom panels). Green box represents the micro-irradiation site and the white arrow represents the line pair correlation analysis is performed across. Scale bar is 3 μm. (c) Schematic of pair correlation analysis of eGFP-53BP1 mobility along a line scan that traverses the laser micro-irradiation site: the time series of frames acquired in (a) contains information on eGFP-53BP1 mobility outside and inside the laser micro-irradiation site in the form fluorescence intensity fluctuations (top and bottom left panels). If we spatially correlate a pair of fluorescence intensity fluctuations at a distance (δr) that enables comparison of eGFP mobility outside the DNA damage site (position 1) with eGFP mobility inside the DNA damage site (position 2) for all possible time delays (τ) we obtain a pair correlation profile that describes the average time eGFP-53BP1 molecules take to traverse this distance (bottom right panel). If we plot this time as a function of pixel position along the line scan we obtain a trace of eGFP mobility (top right panel). (c) When we apply the workflow of analysis described in (b) to the data presented in (a) we are able to correlate eGFP-53BP1 localisation (top panel) with eGFP-53BP1 mobility (bottom panel).

**Supplementary Table 1.**
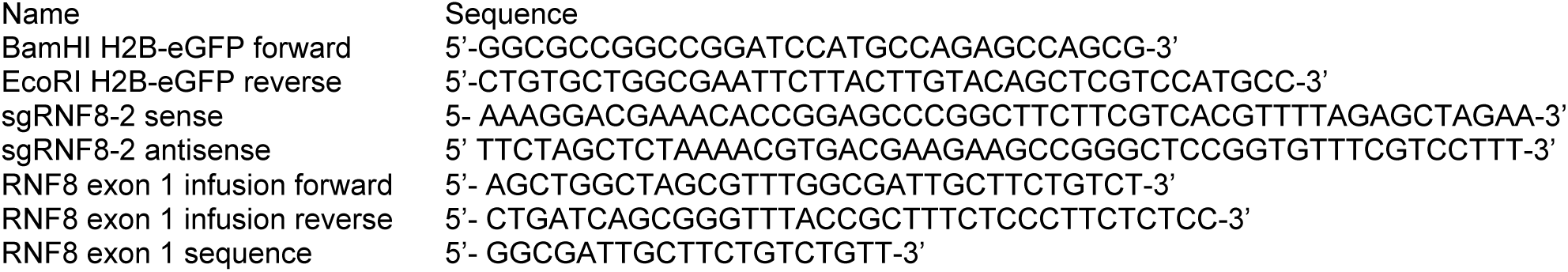
Oligonucleotides used in this study.

